# The acidic N-terminus of HHARI and neddylation are essential for the activation and maintenance of RIG-I-mediated type I interferon response

**DOI:** 10.1101/2025.02.01.636034

**Authors:** Ioanna Kontra, Harry Ward, Faith Vinluan, Rachel Lau, Anna Surace, Vinothini Rajeeve, Pedro Cutillas, Benjamin Stieglitz, Myles J. Lewis

## Abstract

Human homolog of Ariadne (HHARI) is a RING-between-RING ubiquitin E3 ligase which interacts with cullin-RING E3 ligase (CRL) complexes. HHARI has been implicated in the type-I interferon anti-viral response. However, how HHARI drives interferon signalling is not fully understood and the function of the unique, highly conserved acidic N-terminal domain of the protein is unknown. Here, we show that HHARI stimulates interferon-β secretion and autocrine type-I interferon signalling by directly targeting the viral RNA sensor RIG-I (Retinoic Acid-Inducible Gene I) in a neddylation-dependent manner. This suggests that neddylation inhibition could be used to treat interferonopathies and related diseases. Truncated HHARI containing only the N-terminal acidic/UBA-like domains retained the ability to induce interferon signalling in a neddylation-dependent mechanism. HHARI-mediated interferon-β secretion was enhanced by overexpression of cullins 1-5. The N-terminal acidic/UBA-like domain of HHARI is critical for RIG-I activation and interferon signalling, as removal of these domains inactivated the pro-interferon phenotype. We propose a mechanism by which the N-terminus of HHARI interacts with all neddylated cullins leading to endogenous HHARI activation. This suggests a model in which the N-terminus of HHARI ‘unlocks’ and activates neddylated cullins, which in turn are required for activation of HHARI itself. As cullins typically form modular cullin-RING ligase super-assemblies our findings imply that the HHARI N-terminus domain is a critical regulator of the versatile CRL system, which, through widespread protein ubiquitylation, controls many eukaryotic cell functions.

## Introduction

Human homologue of Drosophila Ariadne, HHARI (also known as *ARIH1*), is a RING-in-between-RING (RBR) E3 ligase enzyme in the Ariadne subfamily.^1,2^ RBR E3s are structurally defined by the presence of a C_3_HC_4_-type RING domain, in addition to two conserved Cys/His-rich Zn^2+^-binding domains, IBR and RING2 domains. The cognate E2 enzyme of an RBR binds to the RING1 domain of the enzyme to facilitate the formation of a thioester-linked ubiquitin molecule to a conserved cysteine in the RING2 domain,^3^ after which ubiquitin is conjugated to a residue on the target substrate, which is typically a lysine.

As is common among RBRs,^4–8^ HHARI assumes an autoinhibited conformation whereby the C-terminal Ariadne domain folds over and masks the catalytic cysteine in the RING2 (Rcat) domain. To become active, HHARI undergoes structural rearrangement to allow transfer of ubiquitin to the catalytic cysteine.^5^ There are currently two established mechanisms by which HHARI overcomes autoinhibition: phosphorylation of S427,^9^ found in the Ariadne domain, or interaction with neddylated cullin-RING ligases (CRLs).^10–13^

CRLs are highly versatile, modular protein assemblies responsible for a vast proportion of ubiquitination events across multiple cellular pathways.^14^ CRLs are composed of a cullin protein acting as the primary scaffold, in addition to an N-terminally bound adaptor and substrate receptor subunit which provides the complex’s substrate specificity.^14^ The C-terminus of the complex binds Really Interesting New Gene (RING) containing protein Rbx1/2, which acts as the catalytic module of the CRL and is responsible for both neddylation and ubiquitylation reactions.^15,16^ Neddylation constitutes the conjugation of NEDD8 on the C-terminus of the cullin scaffold and is required for the activation of the CRL complex.^17–19^ Neddylation also allows HHARI to bind CRLs via its UBA-like domain, forming an E3-E3 super assembly which can catalyse ubiquitylation of a diverse range of substrates.^10–12^

Structure determination of HHARI has been limited by difficulties in visualising the N-terminal acidic/glycine-rich domain. Thus, this domain has largely been excluded from structural studies.^5,12^ Despite this, the essential nature of the acidic/glycine-rich domain has been alluded to by N-terminal truncation mutants of HHARI lacking the N-terminus 100 amino acids, which prevented interaction with neddylated cullins in-vivo.^10^

Recently, a direct role of HHARI in the antiviral IFN pathway has been shown in mice with conditional myeloid deficiency of *Arih1*.^20^ These animals were more likely to succumb to HSV-1 infection. Conditional deficiency of *Arih1* rescued the autoimmune lethality associated with *Trex1^-/-^*-related hyperinterferonopathy in mice. The proposed mechanism in this study involved ISGylation of cGAS.^20^ *ARIH1* is an essential gene: homozygous deficiency of *Arih1* in mice is lethal,^21^ and CRISPR deletion of *ARIH1* is lethal to cells in multiple cell lines.^22^

Here we present evidence for HHARI acting as a positive regulator of the viral RNA-sensor RIG-I,^23^ thereby leading to secretion of IFNβ and a type-I IFN specific signature in cells. Our research uses HEK293 cells, which despite being deficient in cGAS protein, ^24^ exhibit a strong interferon signature upon HHARI overexpression. This demonstrates the existence of novel mechanisms of regulation through HHARI which do not rely on cGAS. We report that the N-terminus of HHARI is essential in promoting IFNβ secretion through a neddylation-dependent mechanism. We speculate that the N-terminus of HHARI acts as a ‘key’ for unlocking CRL activity, which can in turn activate HHARI ligase functionality for ubiquitination of HHARI-recruited substrates, such as RIG-I.

## Results

### HHARI overexpression causes a type I IFN signature

We first aimed to determine the effects of overexpression of HHARI on IFN signalling. An ELISA assay was used to measure the quantity of IFNβ in the supernatants from HEK293 cells transiently transfected with FLAG-HHARI or a control vector for 48 hours. HHARI overexpression led to a substantial induction of IFNβ secretion compared to supernatants from control cells (Figure 1a). The secretion of IFNβ due to overexpression of HHARI demonstrated conventional activation of type I IFN signalling, downstream of the IFN alpha receptor (IFNAR), with increased phosphorylation of JAK kinases TYK2 and JAK1, which associate with the cytoplasmic portion of IFNAR subunits 1 and 2 (Figure 1b). Total JAK1 and TYK2 protein levels remained constant after HHARI overexpression (Figure 1b). There was no detectable increase in phosphorylation levels of either JAK2 or JAK3 (Figures 1b & 1c), confirming that this was an IFNAR specific response due to secretion of IFNβ. IFNAR signalling primarily relies on phosphorylation of STAT1 and STAT2, but also causes STAT3 and STAT5 phosphorylation.^25,26^ We observed substantial phosphorylation of STAT1, 2, 3 and 5 (Figure 1d). Thus, HHARI overexpression induces massive secretion of IFNβ leading to autocrine activation of IFNAR and appropriate JAK-STAT activation. Consistently with this, overexpression of kinase dead [JAK1 K908E^27^, TYK2 K930R^28,29^] kinase constructs suppressed HHARI-mediated STAT phosphorylation and ISRE luciferase reporter induction (Figure S1a-b). Treatment of cells with the proteasome inhibitor MG-132 did not block HHARI-mediated STAT phosphorylation (Figure 1d). In line with this, the protein levels of phosphatases (SHP2, PTP-1B, TC-PTP) acting as major negative regulators of JAK-STAT phosphorylation^30^ were unaffected by HHARI overexpression (Figure S1c). Using cytoplasmic and nuclear fractionation, we observed strong nuclear translocation of pSTAT1, 2, 3 and 5 following HHARI overexpression (Figure 1e).

**Figure 1.**
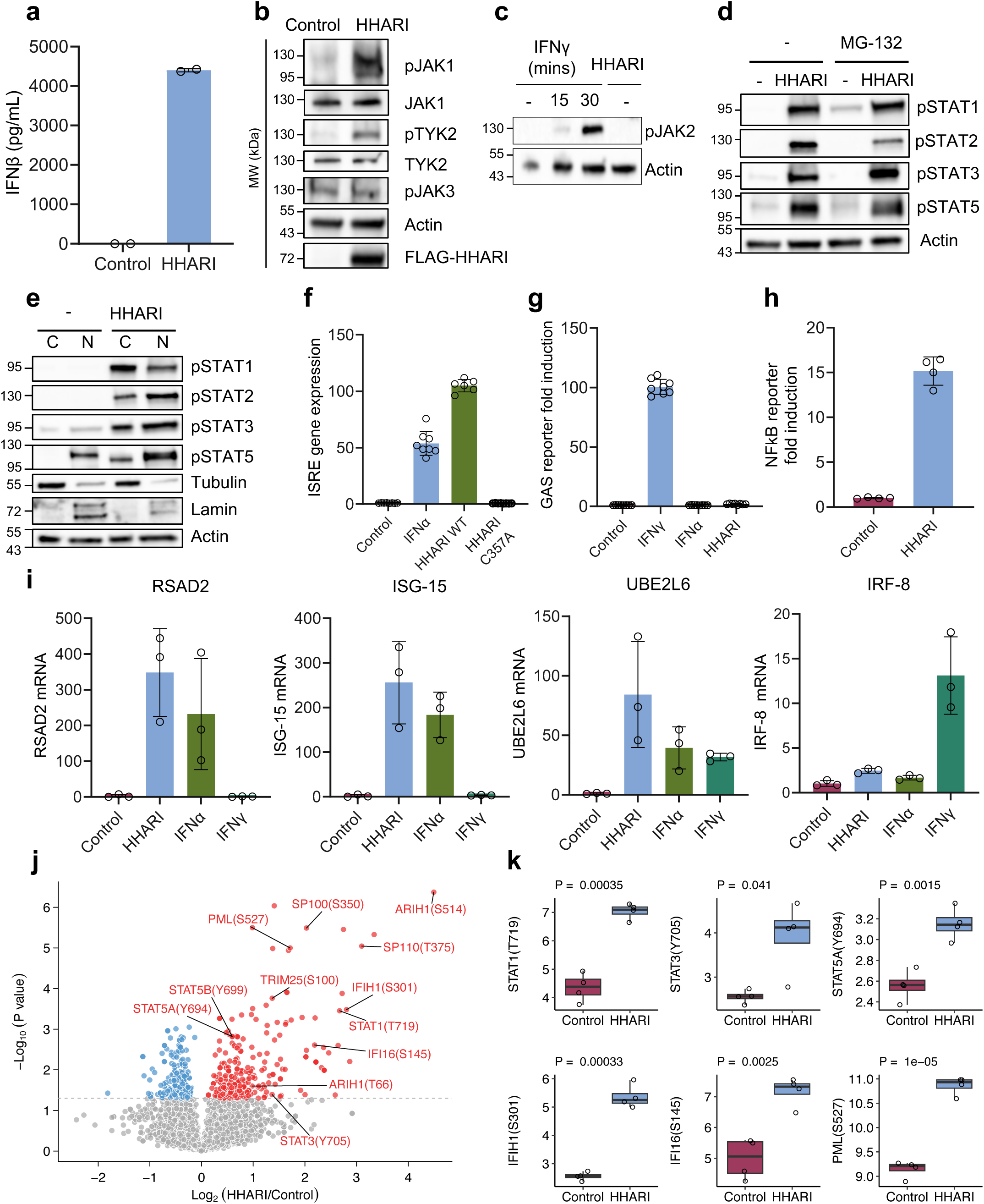
HHARI overexpression leads to a type I IFN signature. **(a)** IFNβ ELISA performed on supernatants from HEK293 cells transiently transfected with FLAG-HHARI or empty vector for 48 hours. **(b-c)** Immunoblot analysis of total and phosphorylated JAK1, TYK2, JAK3 and JAK2 after HHARI overexpression or IFNγ stimulation for the indicated times. **(d)** Cells overexpressing FLAG-HHARI were treated with proteasome inhibitor 10μM MG-132 for 16 hours. Phospho-STAT1, phospho-STAT2, phospho-STAT3 and phospho-STAT5 protein levels were assessed through immunoblot. **(e)** Immunoblot analysis in nuclear (N) or cytoplasmic (C) fractions to assess the protein levels of phospho-STAT1, phospho-STAT2, phospho-STAT3 and phospho-STAT5. Tubulin and Lamin A were used as positive controls for cytosolic and nuclear fractions, respectively. Actin was used as a loading control in all immunoblots. **(f)** ISRE luciferase reporter assay performed on HEK293-Luc2a-GFP-ISRE cells transfected with FLAG-HHARI-WT, -C357A or control vector for 48 hours stimulated with/without 200ng/ml IFNα for 9 hours. Error bars represent the standard deviation. **(g)** GAS luciferase reporter assay on HEK293 Dual IFNγ reporter cells (Invivogen), transfected with FLAG-HHARI or stimulated with 50ng/ml IFNγ/IFNα for 24 hours. **(h)** NFκB luciferase reporter assay on HEK293-NFκB cells transfected with FLAG-HHARI. **(i)** Fold changes in mRNA generated through real time reverse transcriptase PCR are shown. Genes investigated include RSAD2, ISG-15, UBE2L6 and IRF-8. Cells were transfected with FLAG-HHARI or control vector for 48 hours. Cells were stimulated with 100ng/ml IFNα or IFNγ for 24 hours. Error bars represent standard deviation. **(j)** Volcano plot of phosphoproteomics data depicting the fold change of each phosphosite. MatrixTests package (version 0.1.9) was used for t-test analysis. Red circles show phosphosites which have significant increases. Blue circles show phosphosites which have significant decreases. Grey circles are phosphosites without any differences. **(k)** Boxplots showing normalised phosphosite abundance for STAT1 (T719), STAT3 (Y705), STAT5A (Y694), IFIH1 (S301), IFI16 (S145) and PML (S527).

We next examined the gene expression signature associated with the profound activation of JAK-STAT proteins. Although, the combination of JAK1 and TYK2 is more specific to the IFNAR receptor, JAK1 is also utilised by the IFNGR, in combination with JAK2. To determine whether the gene signature in HHARI-transfected cells corresponded to type I IFN ISGs, we compared ISRE (type I IFN specific) and gamma-activated sites (GAS, type II IFN specific) luciferase reporter activation after HHARI overexpression. We found a strong induction of the ISRE reporter by HHARI (Figure 1f), whereas the GAS reporter was only activated by IFNγ stimulation (Figure 1g). This is further evidence towards HHARI activating IFNAR signalling specifically leading to induction of type I interferon-stimulated gene transcription. It is known that NFκB activation is often co-ordinated with transcriptional activation of IFNβ upstream of IFNβ secretion. In line with previous reports,^31,32^ we also observed activation of an NFκB luciferase reporter in cells overexpressing HHARI (Figure 1h). This suggested that HHARI targets a pathway upstream of IRF3 and NFκB activation in order to promote transcription and secretion of IFNβ. In addition to the ISRE and GAS luciferase reporter assays, the expression of specific ISGs was measured through RT-qPCR (Figure 1i). Type-I IFN inducible genes such as *RSAD2* or *ISG15* were confirmed as IFNα-specific and were shown to be upregulated by HHARI (Figure 1i). On the contrary, genes such as *IRF8*, which are more specifically induced by IFNγ, ^33,34^ were not significantly upregulated by HHARI overexpression or IFNα stimulation (Figure 1i). We measured a significant induction of *STAT1* and *STAT2* mRNA after overexpression of HHARI or IFNα/γ stimulation (Figure S1d), consistent with both *STAT1* and *STAT2* being ISGs which are inducible by IFNα/β.^34,35^ Protein levels of selected type-I IFN inducible genes were also verified: both UBE2L6, the key E2 enzyme for ISGylation, and the ISG15 associated deubiquitinase USP18 protein levels were significantly higher in HHARI overexpressing cells (Figure S1e-f).

We also confirmed the presence of a type I IFN-inducible signature through LC-MS/MS (liquid chromatography tandem mass spectrometry) phosphoproteomic analysis of HEK293 cells overexpressing HHARI. A volcano plot was used to visualise significantly differentially regulated phosphosites (Figure 1j). Phosphorylation of STATs 1, 3 and 5 was thus confirmed through this independent technique (Figure 1k). We also observed phosphorylation of a number of IFN-inducible proteins IFIH1, IFI16 and PML in HHARI overexpressing cells compared to control cells (Figure 1k). This confirms that HHARI overexpression leads to a full type I IFN response with induction of interferon stimulated genes at protein level and activation of key ISG proteins as shown by phosphorylation.

### HHARI activates RIG-I to promote IFNβ secretion in a ubiquitination and neddylation-dependent mechanism

We were particularly interested in deciphering the cellular target of HHARI, upstream of IFNβ secretion. Although HEK293 are deficient in cGAS,^24^ we first investigated STING activation, owing to a report that HHARI positively regulates cGAS.^20^ While we were able to observe STING activation in HEK293 cells treated with the STING agonist diABZI, STING was not activated by HHARI overexpression (Figure 2a). To study whether STING inhibition interfered with HHARI-mediated IFNβ secretion, we used the small molecule inhibitor of human STING, H-151. HHARI overexpressing cells treated with H-151 displayed only a slight reduction in IFNβ secretion (Figure 2b) in contrast to the complete inhibition observed with MLN4924 in later experiments (Figure 3). This suggested that in HEK293 cells a STING-independent pathway was the dominant mechanism underlying HHARI-mediated IFNβ secretion.

**Figure 2.**
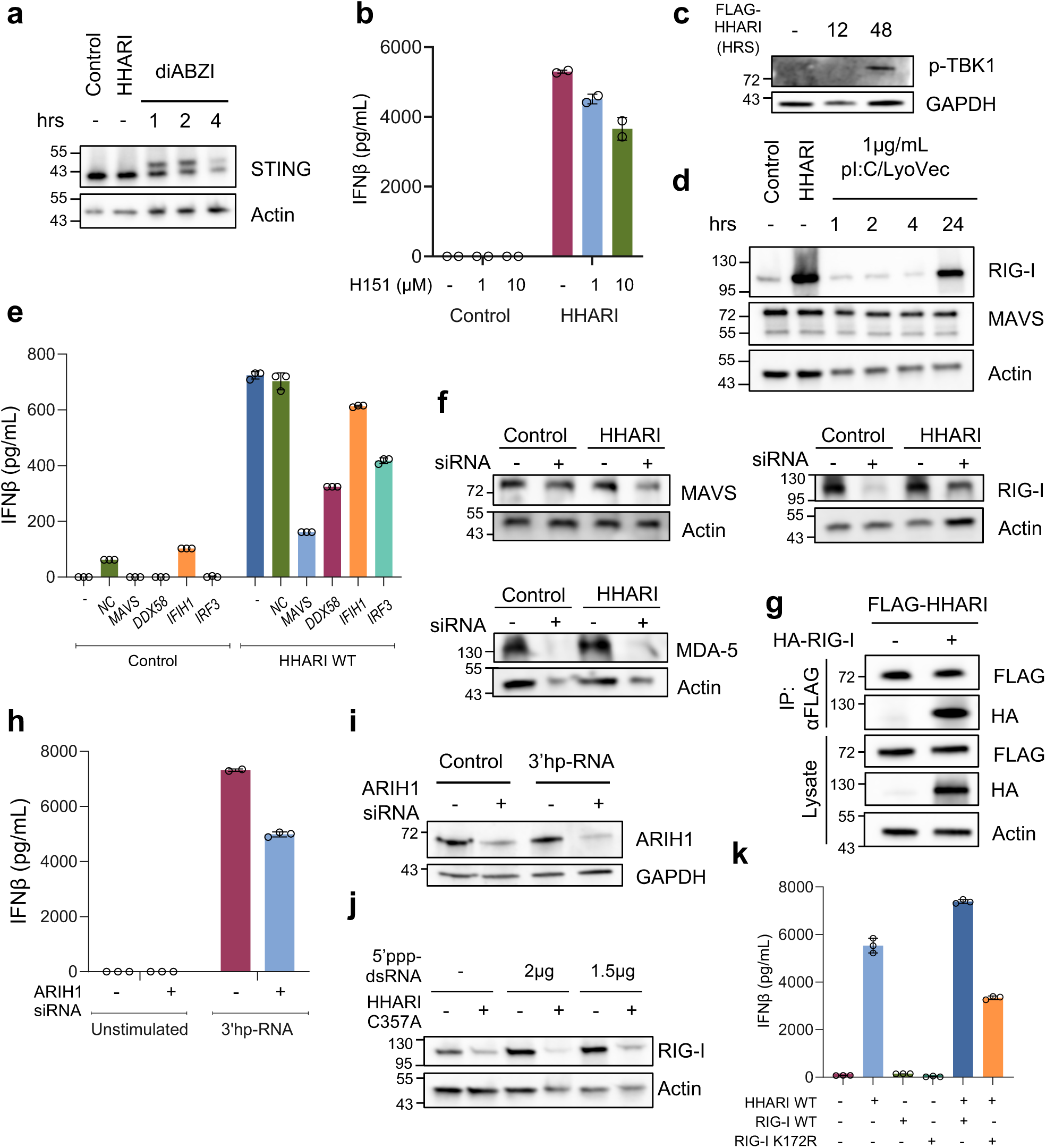
HHARI promotes IFNβ secretion through the RIG-I/MAVS pathway. **(a)** HEK293 cells were transiently transfected with FLAG-HHARI or a control vector. Cells were stimulated with diABZI (STING agonist) for 1, 2 or 4 hours, as a positive control. Immunoblot analysis is shown for STING levels. Actin was used as a loading control. **(b)** IFNβ secretion in cell supernatants was measured from cells transfected with FLAG-HHARI and treated with increasing doses (1-10μM) of human STING inhibitor H-151 for 24 hours. **(c)** Immunoblot analysis for pTBK1 levels in cells overexpressing FLAG-HHARI for 0, 12 or 48 hours. **(d)** Cells were transfected with FLAG-HHARI, a control vector or polyI:C/LyoVec (RLR agonist) for 1, 2, 4 or 24 hours, as a positive control. RIG-I and MAVS activation was measured through immunoblot. **(e)** IFNβ secretion was quantified in supernatants from HEK293 cells co-transfected with FLAG-HHARI and 30nM of siRNA against DDX58 (RIG-I), MAVS, IFIH1 (MDA5) or IRF3. (f) siRNA knockdown efficacy for RIG-I, MAVS and MDA-5 was measured using western blot. Actin was used as a loading control. **(g)** FLAG-HHARI was immunoprecipitated from HEK293 lysates. Interaction with HA-RIG-I was verified via immunoblot on IP eluate. **(h-i)** Cells were transfected with non-targeting or ARIH1-specific siRNA. 48 hours later, cells were stimulated with 1μg/mL 3’hp-RNA for 6 hours. IFNβ ELISA of supernatants (h) and immunoblot of endogenous ARIH1 levels (i) are shown. **(j)** RIG-I activation was measured through immunoblot in HEK293 cells transfected with FLAG-HHARI C357A (ligase defective) with/without 5’ppp-dsRNA (RIG-I agonist) at varying doses. Actin was used as a loading control. **(k)** IFNβ secretion was measured in cells supernatants from cells transfected with FLAG-HHARI with RIG-I WT or K172R mutant.

**Figure 3.**
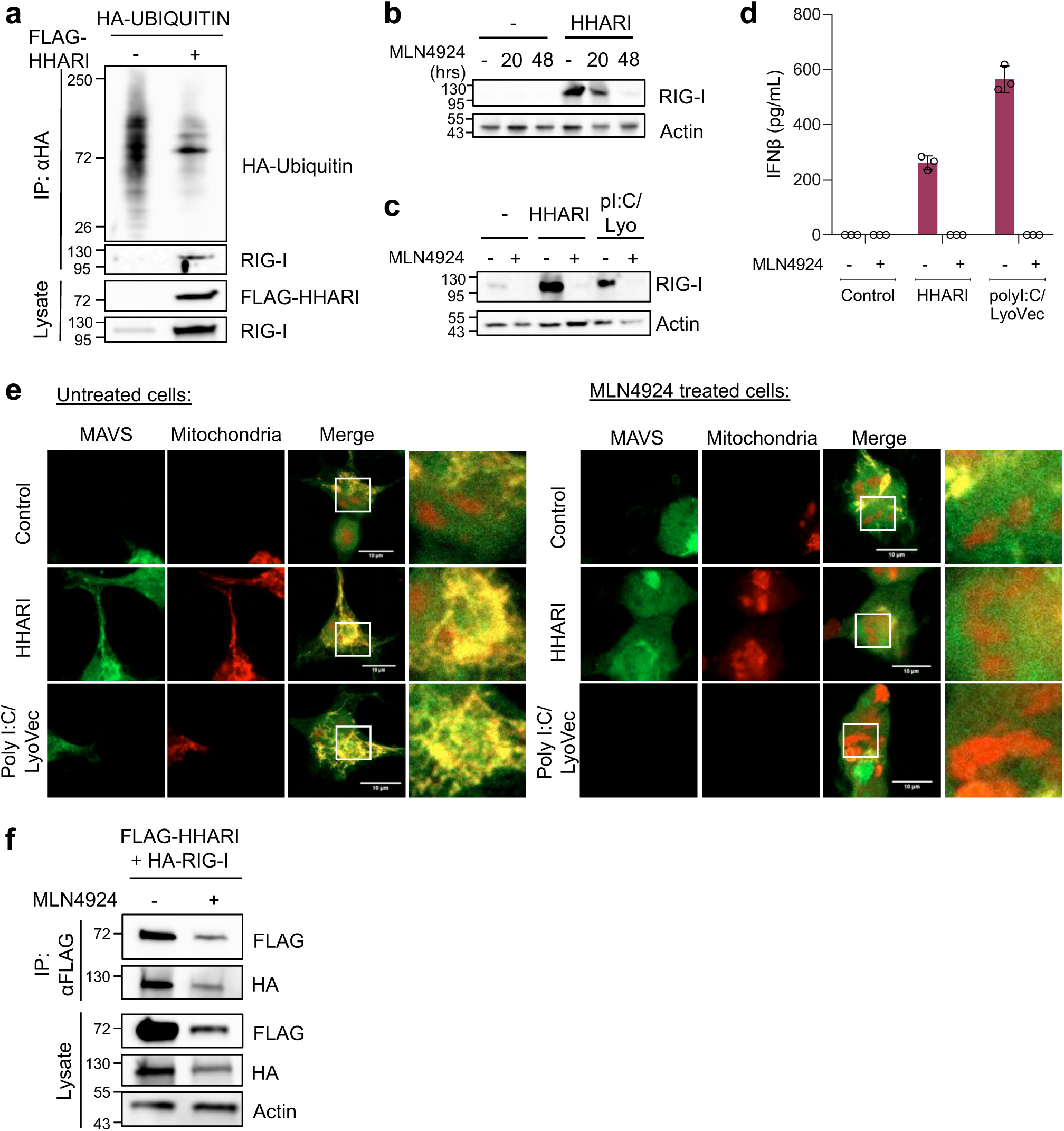
HHARI-mediated activation of the RIG-I/MAVS pathway is ubiquitination and neddylation dependent. **(a)** HA-tagged ubiquitin was immunoprecipitated from HEK293 cells. The presence of ubiquitinated RIG-I was verified in the absence or presence of FLAG-HHARI. **(b-c)** RIG-I induction was measured through immunoblot in cells transfected with FLAG-HHARI or polyI:C/LyoVec (pI:C/Lyo) and treated with 10μM MLN4924 for the time indicated. **(d)** ELISA assay on IFNβ secretion from supernatants of cells treated as described in (c). **(e)** Confocal microscopy assessing MAVS aggregation after FLAG-HHARI or polyI:C/LyoVec transfection. Left panel shows untreated cells whereas right panel shows cells treated with 10µM MLN4924 for 48 hours. HEK293 cells were stained with MitoSpy Orange CMT-MRos (red) and MAVS-Alexa Fluor488 (green). Scale bars represent 10μm. **(f)** FLAG-HHARI was immunoprecipitated from HEK293 lysates. Interaction with HA-RIG-I was assessed in the presence or absence of MLN4924.

We examined TBK1 phosphorylation, a shared kinase amongst STING, RLR and TLR pathways and found elevated phospho-TBK1 levels after HHARI overexpression, in line with its activation (Figure 2c). We investigated RLR pathways and found that HHARI overexpression caused a significant induction of RIG-I protein but not MAVS (Figure 2d). To dissect these pathways further, we studied the effect of siRNA knockdown of RIG-I (*DDX58*), MDA-5 (*IFIH1*, RNA sensor), MAVS (downstream adaptor protein) and IRF3 on HHARI-mediated IFNβ secretion (Figure 2e-f). *DDX58*, *MAVS* and *IRF3* knockdown inhibited the ability of HHARI to promote IFNβ secretion, whereas *IFIH1* siRNA did not have a potent effect (Figure 2e). To probe for a direct interaction between HHARI and RIG-I, we immunoprecipitated FLAG-HHARI from cells overexpressing FLAG-HHARI with/without HA-RIG-I. We detected HA-RIG-I in the FLAG-immunoprecipitated fraction of cells (Figure 2g), which suggests that HHARI interacts with RIG-I to promote its activation. To evaluate HHARI’s significance in RIG-I mediated signalling, we used siRNA to knockdown endogenous HHARI and compared cells’ ability to raise an IFN response when treated with to 3’hp-RNA (a synthetic RNA agonist) (Figure 2h-i). We found that cells containing less endogenous HHARI produced less IFNβ in response to 3’hp-RNA treatment (Figure 2h), re-iterating HHARI’s importance in efficient RIG-I signalling. We also examined whether ligase-defective HHARI-C357A mutant could act as a dominant-negative in cells treated with another RIG-I agonist (5’ppp-dsRNA). Co-transfection of 5’ppp-dsRNA with HHARI-C357A led to the loss of RIG-I induction (Figure 2j). This suggests that HHARI’s ligase is necessary for effective RIG-I activation in response to RNA stimuli. To understand whether HHARI may regulate RIG-I activity through a known ubiquitination site, we mutated RIG-I to include a single amino acid mutation at K172R.^36^ Combining HHARI with WT RIG-I overexpression led to an increase in IFNβ secretion by cells, whereas the RIG-I K172R mutant diminished IFNβ secretion compared to cells overexpressing HHARI only or HHARI and RIG-I WT (Figure 2k).

To confirm whether overexpression of HHARI activates RIG-I through a ubiquitin-mediated mechanism, we overexpressed HA-tagged ubiquitin in HEK293 cells in combination with FLAG-HHARI or a control vector. We immunoprecipitated HA-Ubiquitin in order to enrich for ubiquitinated proteins in control and HHARI-overexpressing cells (Figure 3a). We observed that ubiquitinated RIG-I could be found in HHARI overexpressing cells but not control cells (Figure 3a). Despite the fact that total RIG-I protein increases in the presence of HHARI due to its function as an ISG, research has shown that newly synthesised RIG-I is not subject to mitochondrial aggregation^37^ and thus is unlikely to be ubiquitinated. Hence, the ubiquitinated RIG-I shown in Figure 3a is likely a product of HHARI-mediated activation upstream of the initial release of IFNβ.

Other than the more recent findings of phosphorylation-mediated activation of HHARI’s ligase,^9^ the primary mechanism of activation is considered to be through binding to a neddylated CRL, which in turn displaces HHARI’s Ariadne domain. In order to study the potential involvement of neddylation in HHARI-mediated RIG-I activation, we used the neddylation inhibitor MLN4924. MLN4924 inhibits the NEDD8-activating enzyme (NAE),^38,39^ thus blocking all cullin neddylation and interfering with HHARI-CRL interactions. We found that MLN4924 treatment potently inhibited HHARI-mediated RIG-I activation in a time-dependent manner (Figure 3b). In addition, neddylation inhibition was also detrimental to polyI:C/LyoVec (synthetic RLR agonist) mediated RIG-I activation, causing a reduction in RIG-I protein induction (Figure 3c) and IFNβ secretion (Figure 3d). RIG-I signalling relies on the downstream mitochondrial adaptor protein MAVS for induction of IFNβ. We sought to visualise whether HHARI overexpression or polyI:C/LyoVec stimulation of cells caused changes in MAVS aggregation through confocal microscopy. Cells were stained with MitoSpy Orange CMTMRos (red), fixed, permeabilised and stained with anti-MAVS-Alexa488 antibody (green) and imaged by confocal microscopy. Figure 3e shows representative images from untreated (left panel) or MLN4924-treated (right panel) cells. Increased aggregation and co-localisation of MAVS was observed in HHARI-overexpressing or polyI:C/LyoVec-stimulated cells, consistent with activation of the RLR-MAVS pathway. Treatment of cells with MLN4924 completely suppressed the pattern of MAVS aggregation observed in the untreated cells in both HHARI overexpressing and polyI:C/LyoVec stimulated cells.

We theorised that MLN4924’s inhibition of RIG-I activation and IFNβ secretion may stem from a loss of the HHARI-RIG-I interaction. We immunoprecipitated FLAG-HHARI from untreated cells or cells treated with MLN4924 and examined the amount of co-immunoprecipitated HA-RIG-I (Figure 3f). Although we did observe a reduction, this reduction in HA-RIG-I was proportional to the reduction in the amount of FLAG-HHARI which had been immunoprecipitated. The difference in immunoprecipitation efficiency became apparent when we analysed the whole cell lysate for levels of FLAG-HHARI. The level of recombinant HHARI expression was reduced in MLN4924 treated cells compared to untreated cells. However, HHARI expression in MLN4924 treated cells still exceeded the levels of endogenous HHARI. Despite this, IFNβ secretion was completely suppressed in MLN4924-treated cells overexpressing HHARI (Figure 3d). As a HHARI-RIG-I interaction could still be detected in MLN4924-treated cells, this led us to postulate that neddylation is a prerequisite for the activation of HHARI and therefore a complex of HHARI and RIG-I in the absence of neddylation does not result in RIG-I activation nor IFNβ secretion.

### HHARI N-terminus domain is critical for IFNβ secretion

We proceeded to investigate how the domain structure and individual residues within the HHARI protein might contribute towards its function in RIG-I activation and IFN signalling. Due to the unique nature of HHARI’s N-terminal domain (highly acidic & glycine rich), there is little information available from protein structures which may be indicative of its structural state, molecular interactions and function. We were therefore particularly interested in generating C- and N-terminal domain truncations of HHARI (Figure 4a) in order to investigate how they affect HHARI-mediated IFNβ secretion. We also assayed single amino acid substitution mutants focusing on important catalytic residues or potential phosphorylation sites, as identified by our earlier LC-MS/MS phosphoproteomics study of cells overexpressing HHARI (Figure 4b).

**Figure 4.**
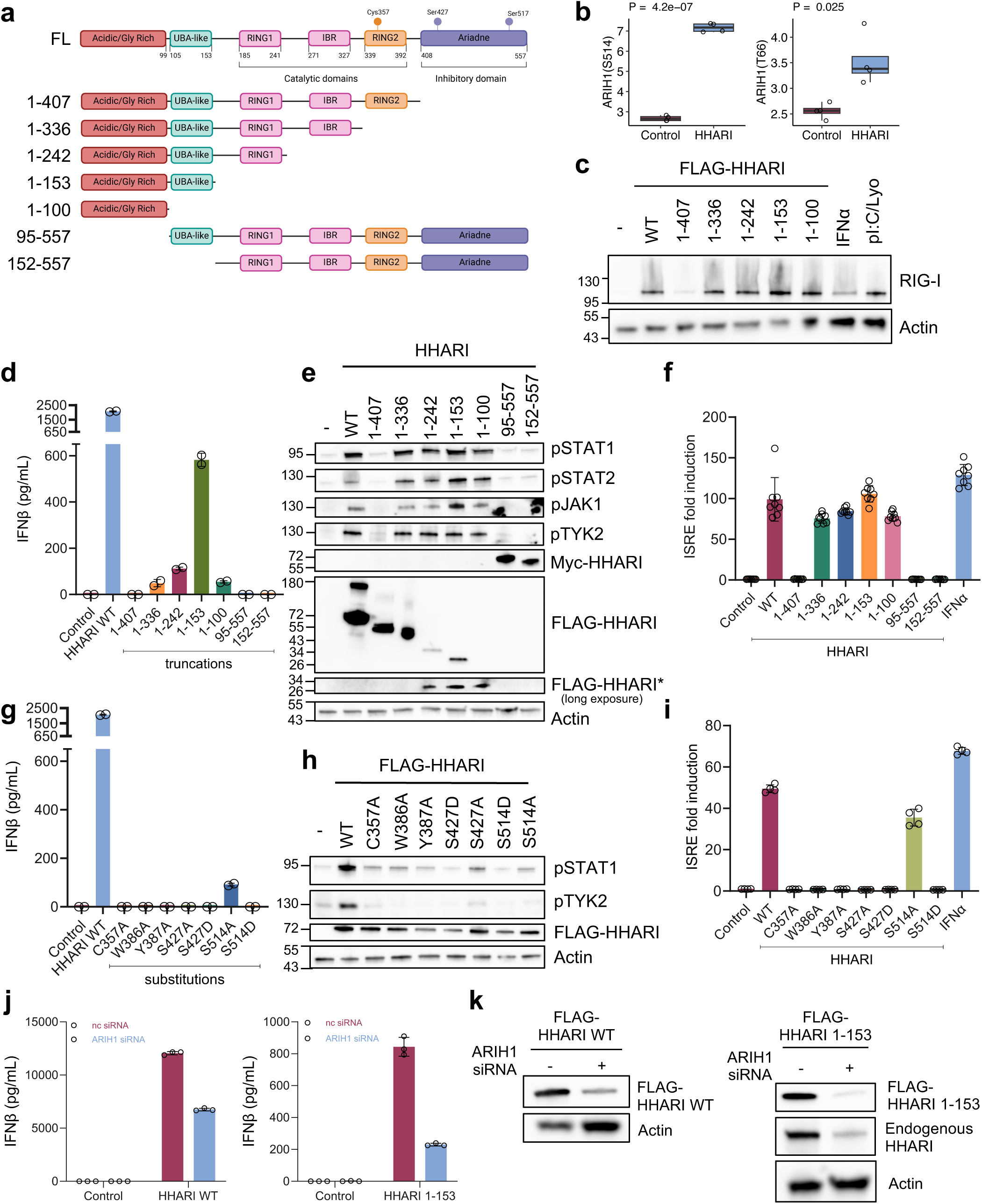
Studying HHARI through site-directed mutagenesis and truncation mutants. **(a)** Domain composition of the full-length (FL) HHARI protein followed by the constructed C- and N-terminus mutants created through site directed mutagenesis or sub-cloning. Cys357 represents the catalytic cysteine. Ser427 and Ser514 represent putative phosphorylation sites. **(b)** Boxplots showing normalised phosphosite abundance for HHARI S514 and HHARI T66. **(c)** Immunoblot for RIG-I protein in cells transfected with FLAG-HHARI WT or truncation mutants. IFNα and polyI:C/LyoVec were used as positive controls. Actin was used as a loading control. **(d-e)** HEK293 WT cells were transiently transfected with FLAG-HHARI WT or truncation constructs, or a control empty vector for 48 hours. Secreted IFNβ was measured in cell supernatants through ELISA assay (d) and IFNAR activation was measured through immunoblot analysis of phopsho-STAT1, phospho-STAT2, phospho-JAK1 and phospho-TYK2. Actin was used as a loading control (e). **(f)** ISRE luciferase reporter assay on supernatants collected from cells transfected with FLAG-HHARI WT or mutant constructs, or a control empty vector for 48 hours. Cells were stimulated with IFNα as a positive control where appropriate. Error bars represent the standard deviation. **(g-i)** HEK293 WT cells were transiently transfected with FLAG-HHARI WT or substitution mutants, or a control empty vector for 48 hours. Supernatants and protein lysates were analysed as described for (d-f), respectively. **(j-k)** Cells were transfected with negative control (nc) or ARIH1-specific siRNA. 48 hours later, HHARI-WT or HHARI 1-153 DNA was transfected for an additional 24 hours. IFNβ ELISA of supernatants is shown (j) as well as immunoblot for FLAG-HHARI and endogenous HHARI protein levels (k). Actin was used as a loading control.

Each HHARI truncation mutant was assayed for its ability to induce RIG-I (Figure 4c), in addition to IFNβ secretion (Figure 4d), JAK-STAT activation (Figure 4e) and ISRE reporter activation (Figure 4f). As the Ariadne domain of HHARI has been linked to autoinhibition of the protein through hinderance of the RBR ligase, we hypothesised that removal of the Ariadne domain (truncation 1-407) may lead to hyper-activation of HHARI and enhance IFNβ secretion. Curiously, removal of the Ariadne domain completely inhibited the ability of HHARI to activate RIG-I, shown by a complete loss of RIG-I induction at the protein level (Figure 4c), as well as loss of detectable IFNβ in cell supernatants (Figure 4d), reduction of JAK-STAT protein phosphorylation to basal levels (Figure 4e) and a complete suppression of ISRE reporter induction (Figure 4f). However, we were surprised by the phenotype presented by the additional truncations following removal of the RING2 domain. HHARI truncations 1-336, 1-242, 1-153 and 1-100 all lack the RING2 domain which contains the catalytic cysteine of the protein required for substrate ubiquitylation. Despite this, each of these mutants, retains the ability to induce RIG-I activation (Figure 4c). We noticed that the 1-153 truncation (acidic/UBA-like domains only) particularly, caused IFNβ secretion (Figure 4d), and saturated IFNAR signalling (Figure 4e) as well as ISRE reporter activation (Figure 4f). Importantly, HHARI 1-153 retains this pro-interferon signature despite being expressed at significantly lower levels compared to the full-length WT HHARI protein (Figure 4e, FLAG blot). The truncation mutants were all expressed at lower levels compared to the full-length protein, with the smallest truncations (1-100, 1-153 and 1-242) showing the most significantly reduced expression with variable expression efficiency (Figure S2a-b). This is due to proteasomal degradation of these truncates (discussed below). N-terminal truncation of the acidic domain (aa 2-94) and UBA-like domains (aa 1-151) led to the inactivation of HHARI. These results suggest that at least the N-terminal 94 amino acids of HHARI are essential for its role in IFN signalling. As expanded upon below, we hypothesise that overexpression of the N-terminal 151 amino acids of HHARI is sufficient to activate endogenous HHARI, whose ligase and full function remains intact.

In addition to domain truncations, the effect of single amino acid substitutions of important catalytic residues in the RING2 domain were also explored. This includes the ligase defective mutant C357A, as well as two substrate recognition deficient mutants W386A and Y387A previously characterised as having a role in substrate binding between HHARI-4EHP.^9^ C357A mutation led to a complete loss of protein activity (Figures 4g-i), suggesting that the ligase activity of HHARI is essential in promoting IFNβ secretion. It is unclear why full-length, catalytically dead HHARI C357A fails to activate endogenous HHARI in the same way the 1-153 fragment does, given that full-length HHARI C357A also possesses the 1-153 region. It is possible that 1-153 and endogenous HHARI can interact in a way that leads to the activation of endogenous HHARI, whereas HHARI C357A may simply block endogenous HHARI. The data suggests that the loss of Ariadne and RING2 domains may allow HHARI 1-153 (as well as aa1-242 and aa1-336 truncations) to activate endogenous HHARI protein, or in other words that the Ariadne and RING2 domains inhibit HHARI C357A from activating endogenous HHARI. Furthermore, W386A and Y387A mutants also led to a loss of activity, suggesting these residues play a role in substrate recognition beyond 4EHP.

A recent study suggested that phosphorylation of the Ariadne domain at S427 bypassed protein autoinhibition and led to increased HHARI ubiquitylation activity.^9^ We were curious to determine whether this was the case *in vivo*. We created phosphomimetic (serine to aspartic acid S427D) and phosphodead (S427A) as well as phosphomimetic (S514D) and phosphodead (S514A) mutations of S514, which was the top hit in the phosphoproteomic mass spectrometry analysis of HHARI-overexpression (Figure 4b). Unexpectedly both phosphomimetic and phosphodead mutation of Ariadne domain residues led to a loss of activity in HHARI (Figures 4g-i). However, it is worth noting that past experiments investigating the role of S427D mutation used *in-vitro* systems and a truncated version of HHARI expressing amino acids 90-557, which may impact on the functionality of Ariadne domain phosphorylation if the two domains associate as a result of protein folding. In addition, evidence of insufficient mimic of phospho-serine by mutation to aspartic acid has been reported in some proteins such as MAVS and IRF3.^40,41^

Data from gnomAD v4.1.0,^42^ which contains whole exome/genome sequencing from 730,947 individuals, shows that predicted loss of function mutations in *ARIH1* are extremely rare in the healthy human population. This strongly reinforces the fact that changes in the amino acid sequence of HHARI are detrimental to protein function and this observation may be linked to the observed cellular phenotype and sensitivity of HHARI for the S427/S514 mutations.

To investigate the relationship between overexpressed HHARI 1-153 and endogenous HHARI protein, we utilised ARIH1-specific siRNA to knockdown endogenous HHARI expression, to assess whether this compromised the 1-153 fragment’s function. As expected, cells treated with ARIH1 siRNA produced less IFNβ when overexpressing HHARI-WT DNA, compared to cells treated with negative control (nc) siRNA (Figures 4j-k). We also found that despite using an siRNA species targeting the C-terminus of HHARI (which the 1-153 fragment lacks), HHARI 1-153’s expression was reduced in the presence of endogenous HHARI knockdown (Figure 4k). This reiterates the hypothesis that there is interplay between endogenous HHARI and the overexpressed 1-153 fragment, suggesting that 1-153 HHARI is capable of interacting with endogenous HHARI protein to promote its functionality.

### HHARI function is enhanced by overexpression of cullins

The acidic and UBA-like domains of HHARI have been implicated in binding neddylated cullins 1-4A.^10^ Substitutions in the NEDD8 binding region at positions V123, I124 and W140 or removal of the N-terminal 100 amino acids interfere with HHARI binding CUL1.^10^ Thus, we investigated whether cullins affected HHARI’s ability to activate RIG-I. We initially hypothesised that a single cullin protein would specifically interact with HHARI and be involved in RIG-I recruitment for ubiquitination leading to IFNβ release. Cullins 1-5 were overexpressed in untreated or MLN4924-treated cells in combination with HHARI WT or 1-153 truncation (Figure 5a-b). None of the cullin proteins alone caused induction of IFNβ secretion, reiterating that HHARI is the rate-limiting protein in this mechanism (Figure 5a). Surprisingly the combination of HHARI with any of cullins 1-5 enhanced the ability of HHARI to secrete IFNβ. Our findings that cullin-4B and cullin-5 enhanced HHARI-mediated IFNβ secretion are especially surprising given that HHARI has only been reported to bind cullins 1-4A.^10^ Cullin-mediated enhancement of IFNβ secretion was entirely dependent on neddylation as it was blocked by MLN4924. Since all tested cullins were able to enhance HHARI’s function, it is conceivable that HHARI requires any neddylated cullin in order to overcome autoinhibition and ubiquitylate RIG-I.

**Figure 5.**
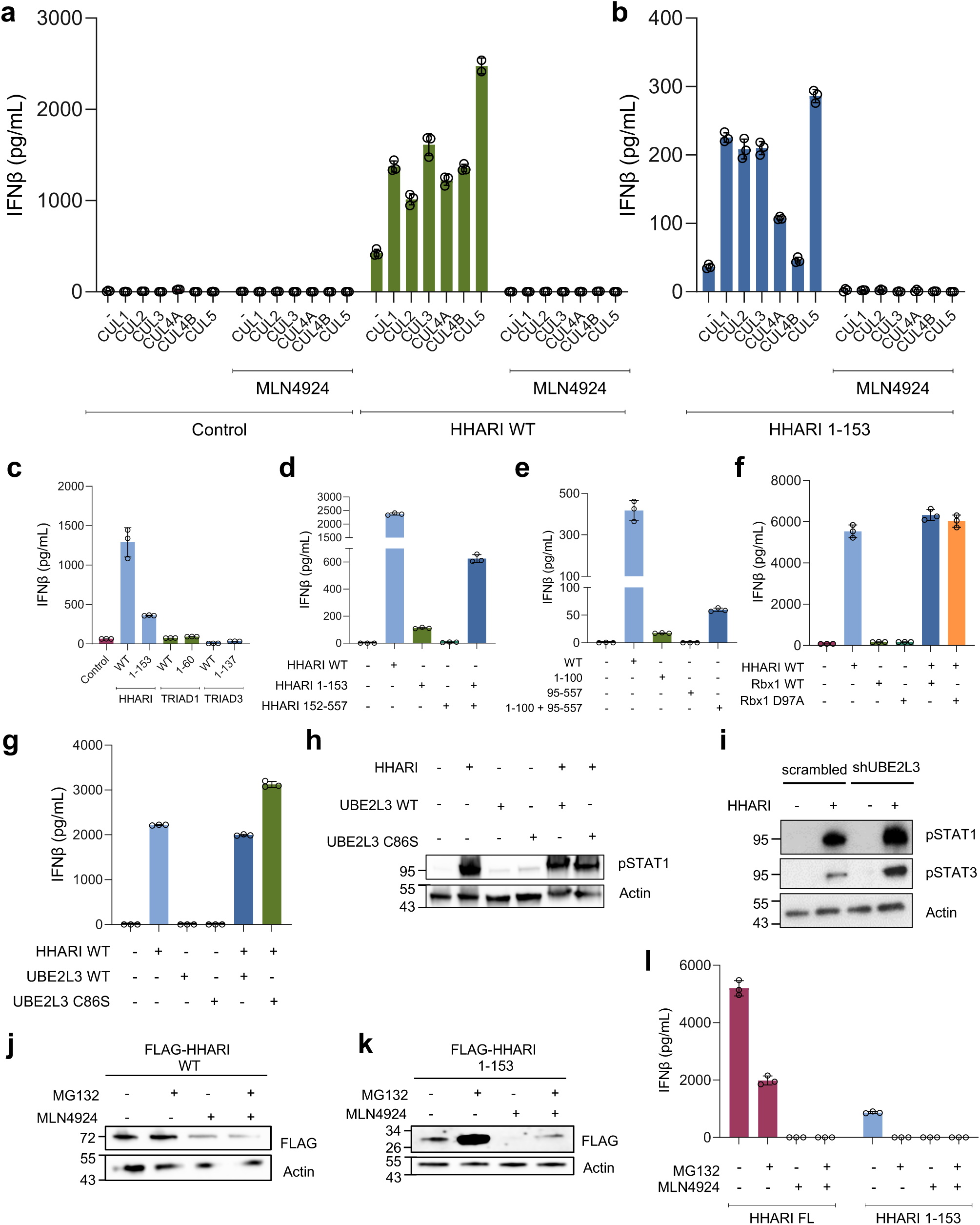
HHARI is activated by cullin overexpression. **(a-b)** IFNβ ELISA on supernatants from HEK293 cells transiently transfected with a control vector, FLAG-HHARI WT or FLAG-HHARI 1-153 with/without myc-CUL1-5 for 48 hours. Appropriate cells were treated with 10μM MLN4924 for 48 hours. Graphs a and b were generated from a single experiment – graphs plotted separately to allow appropriate axis scaling. **(c)** IFNβ ELISA on supernatants from cells overexpressing HHARI, TRIAD1 or TRIAD3 WT or truncated constructs. Error bars represent standard deviation. **(d-e)** IFNβ secretion was measured via ELISA on supernatants from HEK293 cells transfected with FLAG-HHARI WT or a combination of 1-153, 1-100 and/or 153-557, 95-557 truncations for 48 hours. **(f**) IFNβ ELISA on cells overexpressing FLAG-HHARI with/without Rbx1-WT or Rbx1 D97A (ubiquitination deficient, neddylation competent mutant). **(g-h)** HEK293 cells were transiently transfected with a control vector, FLAG-HHARI WT, UBE2L3 WT or UBE2L3-C86S (catalytically dead) for 48 hours. **(g)** Supernatants were collected and assayed for IFNβ via ELISA. **(h)** Immunoblot analysis of phospho-STAT1. **(i)** Scrambled and shUBE2L3 HEK293 cells were transfected with FLAG-HHARI or empty vector for 48 hours. Phospho-STAT1 and phospho-STAT3 protein levels were assessed through immunoblot. Actin was used as a loading control. **(j-l)** Cells were transfected with FLAG-HHARI WT or 1-153 truncation for 48 hours. Cells were treated with 10µM MLN4924 (neddylation inhibitor) for 48 hours and 10μM of proteasome inhibitor MG132 was administered in the final 20 hours of transfection. FLAG-HHARI WT (j) or 1-153 (k) expression was measured through immunoblot. IFNβ secretion was measured via ELISA assay (l).

Based on our findings that the N-terminus of HHARI retains a partial ability to induce IFNβ secretion, we investigated whether this ability is unique to HHARI or shared across other proteins with an acidic N-terminus domain. TRIAD1, a closely related Ariadne RBR ligase (34.9% sequence homology),^43^ and TRIAD3 (also known as RNF216) possess a highly acidic glycine rich N-terminal domain.^44^ We created truncations of TRIAD1 and TRIAD3 containing their respective acidic domains (TRIAD1 aa1-60 and TRIAD3 aa1-137). We found that only HHARI WT and its aa1-153 truncation possessed any significant ability to trigger IFNβ production (Figure 5c). Therefore, this effect is specific to HHARI.

As this truncation lacks the catalytic modules of the protein, we wondered whether it interacted with endogenous HHARI protein in order to promote substrate ubiquitylation. To assess this, we transfected cells with full-length (FL) or truncated mutants of HHARI and measured the activation of IFNβ secretion through ELISA (Figure 5d). We transfected combinations of HHARI 1-153 and HHARI 152-557 to determine if this could reconstitute the effect of the full-length HHARI. We observed diminished induction of IFNβ by HHARI 1-153 compared to HHARI FL, as before. Although HHARI 152-557 did not cause detectable levels of IFNβ release, the combination of 1-153 and 152-557 constructs partially reconstituted the level of IFNβ secretion, although not quite to the level of full-length HHARI. This suggests that the HHARI ligase module is indeed important for promotion of IFNβ secretion in the context of HHARI 1-153. We observed a similar trend of reconstitution, although to a lesser extent, when we combined the aa1-100 truncation of HHARI with the C-terminal 95-557 construct (Figure 5e).

Our findings showing that HHARI’s activity can be enhanced by any cullin left us questioning the relevance of other CRL modules – such as the RING ligase bound to the CTD of cullins (Rbx1/2). Rbx1 is essential for both neddylation of cullins and CRL substrate ubiquitination. Inhibition of neddylation is detrimental to HHARI-mediated IFNβ secretion, so we sought to eliminate Rbx1’s ubiquitin ligase function without hindering its ability to neddylate cullin proteins. To do this, we created a D97A mutant of Rbx1, which is deficient in ubiquitination but neddylation competent.^45,46^ We did not observe any change in IFNβ secretion mediated by overexpression of HHARI when co-expressed with Rbx1 WT/D97A (Figure 5f). This suggests that the ubiquitin ligase activity of Rbx1 is not required for the activation of RIG-I and secretion of IFNβ. Thus, while the neddylated cullin/CRL complex is necessary in order to promote structural re-arrangement and activation of HHARI ligase function, Rbx1 is only required for neddylation and not for ubiquitylation of RIG-I.

### UBE2L3 is not rate-limiting for HHARI-mediated IFNβ secretion

Multiple reports have assigned the cognate E2 enzyme of HHARI as UBE2L3.^2,3^ We examined whether UBE2L3 is rate limiting on HHARI-mediated activation of RIG-I. We observed no significant differences between IFNβ release in cells overexpressing UBE2L3 in combination with HHARI compared to HHARI alone (Figure 5g). We also tested the catalytically inactive UBE2L3 C86S mutant, which is defective in ubiquitin transfer. We observed no reduction in the amount of HHARI-mediated IFNβ release in the presence of UBE2L3 C86S, instead we noted a slight increase in IFNβ production (Figure 5g). A similar trend was observed for the downstream phosphorylation of STAT1 (Figure 5h). STAT1 and STAT3 phosphorylation levels were also increased in shUBE2L3 HEK293 cells overexpressing HHARI compared to the control (scrambled) cell line (Figure 5i). Thus, UBE2L3 is not rate limiting for HHARI’s catalytic activity in the context of RIG-I. HHARI is capable of catalysing in-vitro ubiquitylation in combination with UBE2D1-2 (UbcH5a-b).^47–50^ Hence, these E2 ubiquitin-conjugating enzymes may be relevant in the mechanism by which HHARI promotes the production of IFNβ.

### HHARI expression is destabilised by MLN4924

As we observed a reduction in FLAG-HHARI expression in the presence of MLN4924 (Figure 3e), we sought to investigate whether this was proteasomally mediated by using the proteasome inhibitor MG132 in combination with MLN4924 (Figure 5j). We observed that full-length HHARI protein is not under proteasomal control when overexpressed and the MLN4924-mediated reduction in FLAG-HHARI could not be rescued by inhibition of the proteasome (Figure 5j). This suggests that neddylation inhibition destabilises HHARI expression through other means, possibly endosomal-lysosomal degradation. We were also curious regarding the low levels of expression we observed for HHARI 1-153. We found that basal expression of 1-153 could be significantly increased after proteasomal inhibition using MG132 (Figure 5k), while MLN4924-mediated inhibition of HHARI 1-153 expression was partially rescued by MG132. However, the increase in FLAG-HHARI 1-153 expression did not correlate with an increase in IFNβ release (Figure 5l), suggesting that a functional proteasome is important at some step in the production of IFNβ. However, this could involve another ubiquitin ligase up/downstream of IRF3; or relate to NFκB activation, which requires proteasomal degradation of IκBα, since NFκB activation co-regulates and is required for IFNβ secretion.^31,32^

### HHARI and neddylation promote an interferon-receptor mediated positive feedback loop

Once IFNβ is secreted, it binds to the IFNAR which via a signalling cascade leads to the formation of the trimeric transcription factor ISGF3 (STAT1-STAT2-IRF9), inducing transcription of a large number of anti-viral genes as part of a positive feedback loop.^51^ RIG-I is an important example of this, as it has relatively low expression under basal conditions, but is induced following IFNα receptor stimulation,^34^ contributing to the cell being in an enhanced state of alertness. As part of this positive feedback loop, *IFNB1* and *IFNA* transcription is also increased in an IFNAR-dependent mechanism.^52^ To determine the contribution of the positive feedback loop in HHARI-mediated IFN production, we sought to block IFNAR-mediated signalling through the JAK1/JAK2 inhibitor (Jakinib) Ruxolitinib. We performed a time course of HHARI overexpression ranging from 1 hour to 48 hours after transfection (Figure 6a). IFNβ could be detected in supernatants from 24 hours post-transfection, consistent with the expression of FLAG-HHARI as seen on immunoblot (Figure 6b). We found that IFNβ Cells treated with Ruxolitinib retained a decreased but substantial IFNβ release signature following HHARI overexpression (Figure 6a). Ruxolitinib treatment also prevented the accumulation of IFNβ between the 24 and 48 hour timepoints (Figure 6a), whereas control cells show substantially higher IFNβ secretion at 48 hours, likely attributed to the action of a positive feedback loop. To confirm the lack of IFNAR-mediated signalling, STAT1 phosphorylation was measured (Figure 6c). We observed a complete suppression of STAT1 phosphorylation following treatment with Ruxolitinib, confirming the lack of IFNAR signalling. We also sought to assess whether neddylation regulated this positive feedback loop. Strikingly, we observed that MLN4924 caused a complete loss of ISRE luciferase reporter activation by both HHARI overexpression and direct IFNα stimulation (Figure 6d). The ability of MLN4924 to potently block IFNα-stimulated ISRE reporter induction, strongly suggests that neddylation is additionally required for efficient signalling downstream of IFNAR activation. To confirm whether neddylation is important downstream of IFNAR activation, we stimulated cells with IFNα for 30 minutes (Figure 6e) or overnight (Figure 6f). In both cases, cells were pretreated with MLN4924. STAT1 phosphorylation induced by direct IFNα stimulation was not inhibited by addition of MLN4924, suggesting that neddylation is important downstream of JAK-STAT phosphorylation.

**Figure 6.**
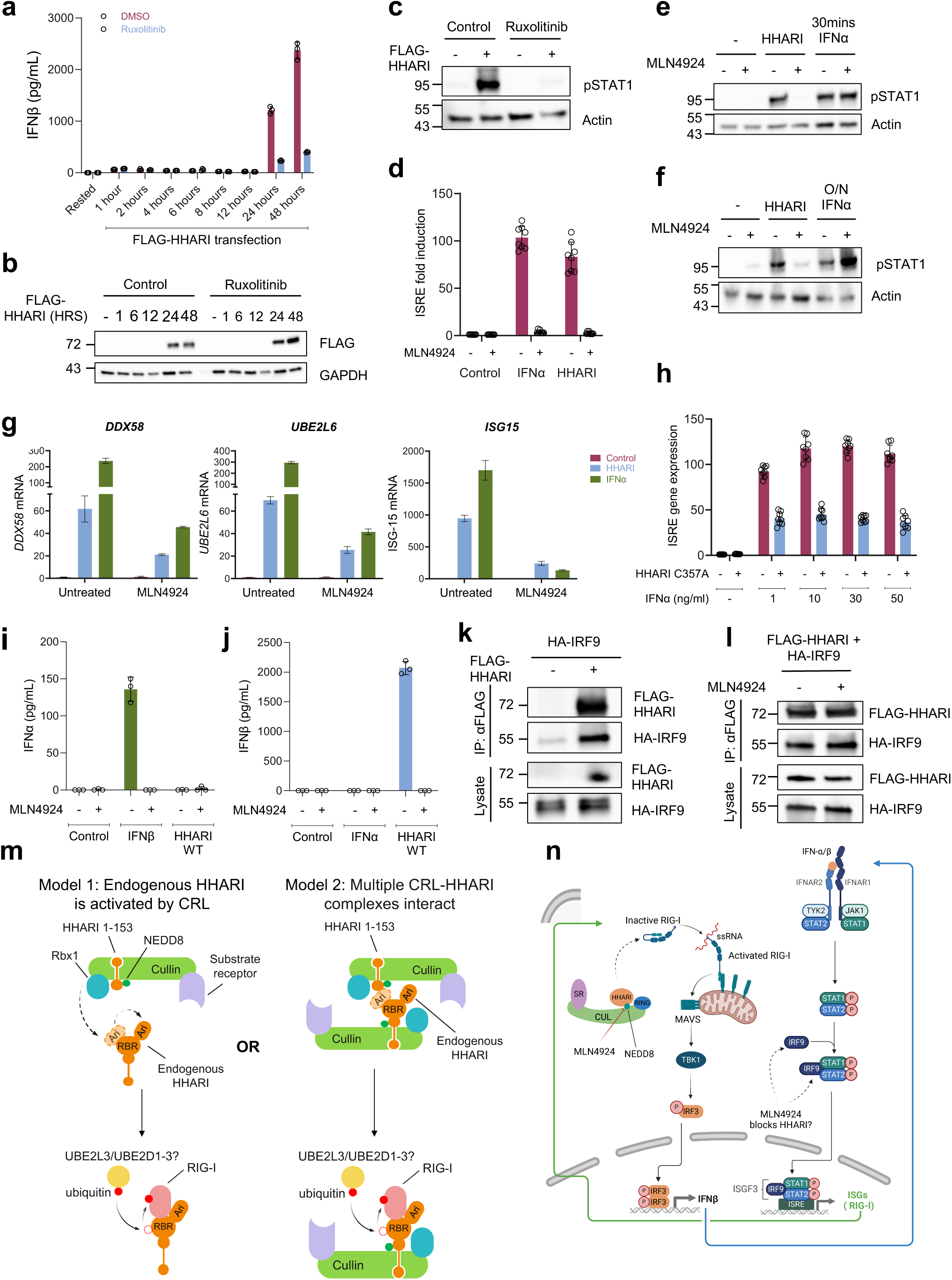
HHARI promotes activation of a neddylation-dependent positive feedback loop downstream of IFNAR signalling. **(a-c)** IFNβ ELISA (a) and western blot (b-c) analysis of cells overexpressing HHARI and treated with JAK inhibitor Ruxolitinib. (b) shows FLAG-HHARI expression and (c) represents pSTAT1 abundance. **(d)** ISRE luciferase reporter assay on HEK293-Luc2a-GFP-ISRE cells transfected with FLAG-HHARI WT or control vector for 48 hours. Appropriate cells were stimulated with 200ng/ml IFNα for 9 hours. Cells were treated with 10µM MLN4924 overnight. Error bars represent the standard deviation. **(e-f)** Cells were transiently transfected with FLAG-HHARI WT or stimulated with 10ng/ml IFNα for 30 minutes (e) or overnight (f). Cells were pre-treated with 10μM MLN4924 for 48 hours. STAT1 phosphorylation was measured through immunoblot. Actin was used as a loading control. **(g)** Fold changes in ISG-15, UBE2L6 and DDX58 mRNA generated through real time reverse transcriptase PCR are shown. Cells were transfected with FLAG-HHARI or control vector for 48 hours or stimulated with 100ng/ml IFNα for 24 hours. Cells were treated with 10μM MLN4924 overnight. Error bars represent standard deviation. **(h)** ISRE luciferase reporter assay on HEK293-Luc2a-GFP-ISRE cells transfected with FLAG-HHARI-C357A (ligase defective) or control vector for 48 hours. Appropriate cells were stimulated with IFNα (1-50ng/ml) for 9 hours. **(i-j)** IFNα (i) and IFNβ (j) ELISA on supernatants from cells stimulated with 100ng/mL IFNβ or 50ng/mL IFNα or transfected with FLAG-HHARI WT, with or without MLN4924 treatment. **(k)** FLAG-HHARI was immunoprecipitated from HEK293 lysates. Interaction with HA-IRF9 was verified via immunoblot on IP eluate. **(l)** FLAG-HHARI was immunoprecipitated from lysates with or without MLN4924 treatment to verify interaction with HA-IRF9. **(m)** Mechanistic diagram depicting the roles of HHARI and neddylation upstream and downstream of IFNβ secretion. **(n)** Proposed mechanism of action for HHARI N-terminal fragment 1-153 through binding to neddylated CRLs, which could promote the activation of endogenous HHARI, leading to ubiquitination of RIG-I.

RT-qPCR analysis of ISG transcripts *DDX58*, *UBE2L6* and *ISG15* revealed that inhibition of neddylation through MLN4924 caused a significant decrease in mRNA induction stimulated by both IFNα and HHARI (Figure 6g). Additionally, we showed that HHARI C357A (ligase defective) strongly inhibits IFNα-mediated ISRE reporter induction (Figure 6h). This suggests that HHARI, as well as neddylation, both play a role in regulating the signalling pathway downstream of IFNAR activation. We also aimed to study whether IFNβ stimulation stimulates further type I IFN production as part of the positive feedback loop. We found that a high dose of IFNβ (100ng/mL) induced secondary IFNα secretion after 48 hours, despite the fact that IFNβ has been documented as acting as a priming agent for response to IFNα stimulation.^53,54^ Thus, this particular effect may be specific to the HEK293 cell system used in this study. HHARI overexpression did not produce any detectable IFNα in cell supernatants (Figure 6i). The effect of IFNβ was neddylation dependent, shown by the loss of IFNα secretion after treatment with MLN4924 (Figure 6i). Thus, this is further evidence of a self-potentiating positive feedback loop which is reliant on neddylation in order to function. We also examined whether IFNα stimulation produced any detectable IFNβ in cell supernatants but could not measure any IFNβ secretion by cells stimulated with IFNα for 48 hours (Figure 6j). A direct interaction between HHARI and IRF9 (a component of the trimer ISGF3) has been previously reported by the BioPlex interactome study.^55^ We were able to confirm this interaction through co-immunoprecipitation (Figure 6k) and show evidence that MLN4924 administration does not interfere with HHARI-IRF9 binding (Figure 6l). Thus, we suggest that HHARI promotes an IFNβ positive feedback loop by interacting with IRF9.

## Discussion

We provide evidence for HHARI and neddylation acting as positive regulators of RIG-I, thereby promoting IFNβ secretion and signalling through IFNAR activation. Dissection of the role of individual HHARI domains revealed that the N-terminal domains of HHARI are essential for its function in the context of IFNβ signalling. Truncation mutants lacking the first 94 or 152 amino acids (HHARI’s acidic/glycine-rich and UBA-like domains) fail to induce IFNβ secretion, an ability which was surprisingly retained by the N-terminal fragment aa1-153, even though this lacks the RBR E3 ligase module. And yet, point mutation of the HHARI catalytic cysteine C357A, shows that the ligase too is essential in the activation of RIG-I. Equally, inhibition of neddylation prevents HHARI-mediated RIG-I activation and IFNβ secretion despite the persistence of HHARI-RIG-I complex formation. HHARI-induced IFNβ secretion was enhanced by overexpression of any of cullins 1-5. We and others^10^ speculate that the acidic N-terminal aa1-153 stretch of HHARI binds to the cullin basic canyon – a highly conserved region within the CTD,^56^ explaining the lack of specificity for a particular cullin. UBE2R1 (Cdc34) is an E2 enzyme with a highly acidic tail, comparable to HHARI’s N-terminus, which nestles in the basic canyon of CUL1.^56^ The acidic tail enables rapid cycling of Cdc34 in and out of the CRL. The structure of UBE2R1’s acidic tail could not be determined until it was co-purified and expressed with a neddylated CRL2^VHL^ complex.^57^ It is conceivable that the structure of HHARI’s N-terminus will not be solved unless co-purified with a neddylated CRL complex. Figure 6m represents various potential modes of action between HHARI and CRLs which may result in activation and RIG-I ubiquitination. One hypothesis is that a CRL complex bound to HHARI 1-153 may activate ‘free’ full-length endogenous HHARI protein (Figure 6m, model 1). Another plausible mechanism could be that the CRL complex bound to HHARI 1-153 interacts with a second CRL complex, which is bound to endogenous HHARI (Figure 6m, model 2). This mechanism could involve the formation of a multimeric structure consisting of several CRL-HHARI bound complexes, such as recently described.^58,59^ Another possible hypothesis is that binding of HHARI 1-153 to a CRL complex causes a state shift (either conformational or covalent modification e.g. ubiquitination/phosphorylation) in the CRL which subsequently enables displacement of 1-153 and binding and activation of full-length endogenous HHARI, thereby facilitating ubiquitination of RIG-I. Currently available X-ray derived structures of HHARI-SCF complexes do not include the N-terminus domain of HHARI due to difficulties in visualisation using the full-length HHARI protein.^5,12^ Future structural studies into the precise binding modality between HHARI and neddylated CRLs are required to understand the functional role of the acidic domain. There is clear synergy between HHARI and CRLs. The function of HHARI is dependent on, and rate limited by a neddylated (and therefore activated) CRL. Equally, the HHARI RBR ligase has previously been implicated in levels of CRL neddylation, although the mechanism by which this occurs is not understood.^10^ Dysfunctional HHARI ligase with the C357S mutation has also been shown to impair CRL substrate turnover in-vivo.^11^ We believe that in addition to the ligase, the N-terminus of HHARI also regulates CRL activity by promoting ‘active’ complex conformations, with this domain contributing to reciprocal regulation between HHARI and CRLs.

Our data also suggest a further role beyond autoinhibition for the Ariadne domain of HHARI. A previous study has suggested that this domain may also be involved in substrate recognition.^11^ Our data show that point mutations in the Ariadne domain abolish IFNβ secretion, suggesting that these phosphorylation sites are essential, and paradoxically so does truncation of the autoinhibitory Ariadne domain. We believe that this can be explained by the nature of interactions between RIG-I, HHARI and CRLs. We suggest that RIG-I binds to the RING2 domain, as has previously been described for 4EHP,^9^ and involves the 386 and 387 residues as evidenced by the W386A and Y387A mutants (Figure 4f-h). The 1-407 HHARI truncation lacks the essential phosphorylation sites found on the Ariadne domain but retains the ability to bind to RIG-I and CRLs. As a result, this truncation acts as a dominant negative against endogenous HHARI by competing for RIG-I binding but being unable to ubiquitylate it, thus resulting in HHARI 1-407 being unable to induce IFNβ production. In contrast, smaller truncations lacking the RING2 cannot bind RIG-I and act through binding to and activating endogenous neddylated CRLs as detailed above.

HHARI and neddylation additionally function at a secondary point to potentiate RIG-I activation through the maintenance of a positive feedback loop involving ISRE-stimulated gene expression. A simplified schematic representation of HHARI’s function upstream and downstream of IFNβ secretion is shown in Figure 6n. Our data suggests that the target is downstream of JAK-STAT phosphorylation, as MLN4924 treatment did not inhibit IFNα-mediated STAT1 phosphorylation (Figure 6e-f), but did block the ISRE reporter (Figure 6d). We (as well as the BioPlex interactome study) found a direct interaction between HHARI and IRF9 by co-immunoprecipitation (Figure 6k).^55^ Thus HHARI not only initiates but also propagates positive feedback of IFNβ secretion, explaining the level of potency of HHARI as an activator of IFNβ.

Our proposed mechanism is novel in various aspects. We use functional read-outs *in-vivo* to give true functional insights into structural modifications in HHARI and CRL interactions. We propose an essential function of the acidic aa1-153 stretch, which has often been overlooked in structural studies, for binding to any cullin. We also show that HHARI directly binds its substrate RIG-I, rather than through substrate adaptors on a specific cullin, and that HHARI ligase alone is essential for this mechanism rather than through co-operation with the Rbx1 ubiquitin ligase. HHARI not only mediates initiation of IFNβ secretion through RIG-I activation but also modulates autocrine positive feedback downstream of the IFNAR.

Our study has clinical translation: MLN4924, also called pevonedistat, is in phase 2/3 clinical trials for treatment of multiple haematological and solid cancers.^60^ Our data suggest that neddylation inhibition could be used to treat interferonopathies^61^ and related diseases with interferon dysregulation such as systemic lupus erythematosus (SLE).^62^

## Materials and Methods

### Cell Culture, plasmids and transfection

HEK293 cells (ATCC) were maintained in DMEM supplemented with 10% FBS. Cells were transiently transfected with different combinations of plasmids for 48 hours using Fugene HD (Promega). DNA-siRNA co-transfections were performed using Jetprime transfection reagent according to the manufacturer’s instructions. A list of DNA vectors and siRNA oligos used in this work can be found in Supplementary Table 1.

### Site directed mutagenesis and sub-cloning

Several mutant HHARI constructs were generated using site directed mutagenesis or subcloning (as indicated in Table S1). All mutagenesis primers were HPLC purified and synthesised from ThermoFisher Scientific. The PCR reaction was set up on a GeneAmp 9700 thermocycler according to the manufacturer’s protocol (Quikchange II XL, Agilent Technologies). All generated mutants were sequenced using sanger sequencing. N-terminal truncated mutants of HHARI were created using PCR subcloning with restriction enzymes MluI and HindIII. The New England Biolabs (E5000S) Taq PCR kit was used to amplify the desired HHARI ORF for subsequent ligation into pCMV6-Myc (Origene).

### Immunoprecipitation, Immunoblotting and antibodies

Cells were lysed in RIPA buffer containing protease (ThermoScientific, A32953) and phosphatase (Roche, 4906837001) inhibitors. Protein samples were quantified using Bradford protein assay and denatured before being separated on 4-20% SDS-PAGE (BIO-RAD, 4561096) and transferred to a nitrocellulose membrane. Membranes were blocked with 5% milk (in 0.1% TBS-T) followed by immunoblotting with the appropriate primary antibody overnight at 4°C with mild agitation. Membranes were incubated with HRP-conjugated antibodies before being subjected to ECL reagents for detection through the Syngene G:Box Chemi XX6/XX9 imager. For immunoprecipitation of target proteins, the Pierce Classic Magnetic IP/Co-IP Kit (ThermoFisher, 88804) was used according to manufacturer’s instructions. Following cell lysis with 1% NP40 buffer, protein concentration was quantified and 1,000µg of protein was combined with 5µg of immunoprecipitation antibody. Immune complexes were formed by incubation overnight at 4°C or at RT for 1 hour. 25µl of magnetic beads were combined with the protein-antibody solution and incubated at room temperature for 1 hour. The beads were collected using a magnetic stand. Flow-through was collected in a clean tube for analysis. Samples were eluted using lane marker sample buffer (0.3M Tris·HCl, 5% SDS, 50% Glycerol, pH 6.8) supplemented with 50mM Dithiothreitol (DTT). Eluate was collected using the magnetic stand and analysed for protein content through western blot as described above. A full list of antibodies used can be found in Supplementary Table 2.

### ELISA assay

The DuoSet human IFN-β kit (R&D systems, DY814-05) was used according to manufacturer’s instructions, to detect IFNβ in the supernatants of transfected or stimulated cells. A FluoStar Omega (BMG Labtech) plate reader was used to measure optical density (OD) at 450nm. Wavelength correction was used based on OD readings a 570nm, to correct for optical imperfections in the plate.

### Luciferase reporter assay

ISRE-Luc2a-GFP HEK293, STAT3-SIE-Luc2p HEK293 or Dual (SEAP/Luciferase) IFNγ HEK293 (Invivogen) cells were transiently transfected with different combinations of plasmids for 48 hours using Fugene HD (Promega) and/or stimulated with IFNα, IFNγ or IL-6. Luciferase activity was assayed using One-Glo (Promega) or QuantiLuc-4 (Invivogen) and normalised to cell viability using Cell-Titre Glo (Promega) measured on a Berthold Orion luminometer.

### RNA extraction and RT-qPCR

RNA was isolated using the RNeasy Plus Micro Kit (Qiagen, 74034) according to manufacturer’s instructions. RNA purity was verified with a Nanodrop spectrophotometer (ThermoFisher Scientific, 2000c) and later reverse-transcribed to cDNA using the Superscript III first-strand synthesis system (Invitrogen, 18080051). cDNA was quantified by real-time quantitative PCR using the Taqman gene expression master mix (Invitrogen, 4369016) and gene specific Taqman reagents on StudioQuant QS5 Real-Time PCR System. All data was normalised to β-Actin levels.

### Confocal microscopy

Transfected HEK293 cells were plated on a 4-well microscope slide (Hendley-Essex, PH005). Cells were allowed to attach for 5 hours before being stained for mitochondria using 500nM of MitoSpy Orange CMTMRos (Biolegend, 424803). Cells were incubated with MitoSpy solution for 30 minutes at 37°C, 5% CO_2_. Cells were fixed with 4% paraformaldehyde (PFA) before proceeding with permeabilization using 0.1% Triton X-100 solution. Wells were blocked for 30 minutes with blocking solution (1% BSA, 22.52mg/mL glycine in PBS-T) in a humidified chamber. Primary antibody against MAVS (Santa cruz biotechnology, sc-166583AF488) diluted 1:50 in blocking buffer was added to each well. Cells were incubated overnight ay 4°C in a humidified chamber, covered from light. The following day cells were mounted with a coverslip and ProLong Diamond antifade mountant (ThermoFisher Scientific, P36965). Coverslips were sealed and allowed to dry at room temperature being imaged. The LSM800 Zeiss microscope was used to image the cells using the ZenBlue software. Images were processed using ImageJ software.

### Phosphoproteome analysis

Phospho-proteomic experiments were performed using mass spectrometry as reported in ^63,64^ and in supplementary material and methods.

## List of Supplementary Materials

Supplementary Materials and Methods

Supplementary Table 1: List of reagents used, including expression vectors and siRNA species.

Supplementary Table 2: List of antibodies used.

Supplementary Figure 1: HHARI overexpression leads to type I IFN signature.

Supplementary Figure 2: Expression of HHARI truncation mutant varies.

Supplementary References

## Acknowledgements

Ioanna Kontra was supported by a PhD fellowship from Barts Charity (G-001922). Harry Ward was supported by an NIHR (National Institute for Health Research) Academic Clinical Fellowship and Barts Charity support grant G-002936. This study was supported by Barts NIHR Biomedical Research Centre. This study was also supported by funds raised by Jim and Sheila Mansell. The HEK293-Luc2a-GFP-ISRE stable reporter cell line was gifted by Dr Helen Rowe. We acknowledge the CMR Advanced Bio-Imaging Facility of QMUL for the use, help and advice with microscopy.

## References

1. Moynihan, T. P. et al. The Ubiquitin-conjugating Enzymes UbcH7 and UbcH8 Interact with RING Finger/IBR Motif-containing Domains of HHARI and H7-AP1*. (1999) doi:10.1074/jbc.274.43.30963.

2. Ardley, H. C., Tan, N. G. S., Rose, S. A., Markham, A. F. & Robinson, P. A. Features of the Parkin/Ariadne-like Ubiquitin Ligase, HHARI, That Regulate Its Interaction with the Ubiquitin-conjugating Enzyme, UbcH7. JBC 276, 19640–19647 (2001).

3. Wenzel, D. M., Lissounov, A., Brzovic, P. S. & Klevit, R. E. UBCH7 reactivity profile reveals parkin and HHARI to be RING/HECT hybrids. Nature 474, 105–108 (2011).

4. Chaugule, V. K. et al. Autoregulation of Parkin activity through its ubiquitin-like domain. EMBO Journal 30, 2853–2867 (2011).

5. Duda, D. M. et al. Structure of HHARI, a RING-IBR-RING ubiquitin ligase: Autoinhibition of an Ariadne-family E3 and insights into ligation mechanism. Structure 21, 1030–1041 (2013).

6. Stieglitz, B., Morris-Davies, A. C., Koliopoulos, M. G., Christodoulou, E. & Rittinger, K. LUBAC synthesizes linear ubiquitin chains via a thioester intermediate. EMBO Rep 13, 840–846 (2012).

7. Trempe, J.-F. et al. Structure of Parkin Reveals Mechanisms for Ubiquitin Ligase Activation. Science (1979) 340, 1451–1455 (2013).

8. Wauer, T. & Komander, D. Structure of the human Parkin ligase domain in an autoinhibited state. EMBO Journal 32, 2099–2112 (2013).

9. Reiter, K. H. et al. Cullin-independent recognition of HHARI substrates by a dynamic RBR catalytic domain. Structure 30, 1269–1284.e6 (2022).

10. Kelsall, I. R. et al. TRIAD1 and HHARI bind to and are activated by distinct neddylated Cullin-RING ligase complexes. EMBO Journal 32, 2848–2860 (2013).

11. Scott, D. C. et al. Two Distinct Types of E3 Ligases Work in Unison to Regulate Substrate Ubiquitylation. Cell 166, 1198–1214.e24 (2016).

12. Horn-Ghetko, D. et al. Ubiquitin ligation to F-box protein targets by SCF–RBR E3–E3 super-assembly. Nature 590, 671–676 (2021).

13. Hill, S. et al. Robust cullin-ring ligase function is established by a multiplicity of polyubiquitylation pathways. Elife 8, (2019).

14. Lydeard, J. R., Schulman, B. A. & Harper, J. W. Building and remodelling Cullin-RING E3 ubiquitin ligases. EMBO Rep 14, 1050–1061 (2013).

15. Huang, D. T. et al. E2-RING Expansion of the NEDD8 Cascade Confers Specificity to Cullin Modification. Mol Cell 33, 483–495 (2009).

16. Zheng, N. et al. Structure of the Cul1-Rbx1-Skp1-F box Skp2 SCF ubiquitin ligase complex. Nature 416, 703–709 (2002).

17. Duda, D. M. et al. Structural Insights into NEDD8 Activation of Cullin-RING Ligases: Conformational Control of Conjugation. Cell 134, 995–1006 (2008).

18. Baek, K., Scott, D. C. & Schulman, B. A. NEDD8 and ubiquitin ligation by cullin-RING E3 ligases. Current Opinion in Structural Biology vol. 67 101–109 Preprint at 10.1016/j.sbi.2020.10.007 (2021).

19. Baek, K. et al. NEDD8 nucleates a multivalent cullin–RING–UBE2D ubiquitin ligation assembly. Nature 578, 461–466 (2020).

20. Xiong, T. C. et al. The E3 ubiquitin ligase ARIH1 promotes antiviral immunity and autoimmunity by inducing mono-ISGylation and oligomerization of cGAS. Nat Commun 13, (2022).

21. Meehan, T. F. et al. Disease model discovery from 3,328 gene knockouts by the International Mouse Phenotyping Consortium. Nat Genet 49, 1231–1238 (2017).

22. Behan, F. M. et al. Prioritization of cancer therapeutic targets using CRISPR–Cas9 screens. Nature 568, 511–516 (2019).

23. Yoneyama, M. et al. The RNA helicase RIG-I has an essential function in double-stranded RNA-induced innate antiviral responses. Nat Immunol 5, 730–737 (2004).

24. Wiser, C., Kim, B., Vincent, J. & Ascano, M. Small molecule inhibition of human cGAS reduces total cGAMP output and cytokine expression in cells. Sci Rep 10, (2020).

25. Tanabe, Y. et al. IMMUNOLOGY Cutting Edge: Role of STAT1, STAT3, and STAT5 in IFN-Responses in T Lymphocytes. (2005).

26. Su, L. & David, M. Distinct Mechanisms of STAT Phosphorylation via the Interferon-/ Receptor SELECTIVE INHIBITION OF STAT3 AND STAT5 BY PICEATANNOL*. J Biol Chem 275, 12661–12666 (2000).

27. Haan, S. et al. Dual Role of the Jak1 FERM and Kinase Domains in Cytokine Receptor Binding and in Stimulation-Dependent Jak Activation 1. The Journal of Immunology 180, 998–1007 (2008).

28. Gauzzi, M. C. et al. Interferon-dependent Activation of Tyk2 Requires Phosphorylation of Positive Regulatory Tyrosines by Another Kinase*. JBC 271, 20495–20500 (1996).

29. Li, Z. et al. Two Rare Disease-Associated Tyk2 Variants Are Catalytically Impaired but Signaling Competent. The Journal of Immunology 190, 2335–2344 (2013).

30. Chen, W., Daines, M. O. & Khurana Hershey, G. K. Turning off signal transducer and activator of transcription (STAT): The negative regulation of STAT signaling. Journal of Allergy and Clinical Immunology vol. 114 476–489 Preprint at 10.1016/j.jaci.2004.06.042 (2004).

31. Honda, K., Takaoka, A. & Taniguchi, T. Type I Inteferon Gene Induction by the Interferon Regulatory Factor Family of Transcription Factors. Immunity 25, 349–360 (2006).

32. Kim, T. K. & Maniatis, T. The Mechanism of Transcriptional Synergy of an In Vitro Assembled Interferon-Enhanceosome. Mol Cell 1, 119–129 (1997).

33. Indraccolo, S. et al. Identification of Genes Selectively Regulated by IFNs in Endothelial Cells 1. The Journal of Immunology 178, 1122–1135 (2007).

34. Liu, S. Y., Sanchez, D. J., Aliyari, R., Lu, S. & Cheng, G. Systematic identification of type I and type II interferon-induced antiviral factors. Proc Natl Acad Sci U S A 109, 4239–4244 (2012).

35. Der, S. D., Zhou, A., Williams, B. R. G. & Silverman, R. H. Identification of genes differentially regulated by interferon, or using oligonucleotide arrays. Medical Sciences Communicated by George R. Stark, Cleveland Clinic Foundation 95, 15623–15628 (1998).

36. Gack, M. U. et al. TRIM25 RING-finger E3 ubiquitin ligase is essential for RIG-I-mediated antiviral activity. Nature 446, 916–920 (2007).

37. Thoresen, D. T., Galls, D., Götte, B., Wang, W. & Pyle, A. M. A rapid RIG-I signaling relay mediates efficient antiviral response. Mol Cell 83, 90–104.e4 (2023).

38. Nawrocki, S. T., Griffin, P., Kelly, K. R. & Carew, J. S. MLN4924: a novel first-in-class inhibitor of NEDD8-activating enzyme for cancer therapy. Expert Opin Investig Drugs 21, 1563–1573 (2012).

39. Soucy, T. A. et al. An inhibitor of NEDD8-activating enzyme as a new approach to treat cancer. Nature 458, 732–736 (2009).

40. Lin, R., Mamane, Y. & Hiscott, J. Structural and Function Analysis of Interferon Regulatory Factor 3: Localisation of the Transactivation and Autoinhibitory Domains. Mol. Cell Biol. 19, 2465–2474 (1999).

41. Liu, S. et al. Phosphorylation of innate immune adaptor proteins MAVS, STING, and TRIF induces IRF3 activation. Science (1979) 347, (2015).

42. Karczewski, K. J. et al. The mutational constraint spectrum quantified from variation in 141,456 humans. Nature 581, 434–443 (2020).

43. Vinci, M. et al. A de novo ARIH2 gene mutation was detected in a patient with autism spectrum disorders and intellectual disability. Sci Rep 14, (2024).

44. Wang, P., Dai, X., Jiang, W., Li, Y. & Wei, W. RBR E3 ubiquitin ligases in tumorigenesis. Seminars in Cancer Biology vol. 67 131–144 Preprint at 10.1016/j.semcancer.2020.05.002 (2020).

45. Chen, A. et al. The Conserved RING-H2 Finger of ROC1 Is Required for Ubiquitin Ligation*. JBC 275, 15432–15439 (2000).

46. Megumi, Y. et al. Multiple roles of Rbx1 in the VBC-Cul2 ubiquitin ligase complex. Genes to Cells 10, 679–691 (2005).

47. Dove, K. K. et al. Structural Studies of HHARI/UbcH7∼Ub Reveal Unique E2∼Ub Conformational Restriction by RBR RING1. Structure 25, 890–900.e5 (2017).

48. Dove, K. K., Stieglitz, B., Duncan, E. D., Rittinger, K. & Klevit, R. E. Molecular insights into RBR E3 ligase ubiquitin transfer mechanisms. EMBO Rep 17, 1221–1235 (2016).

49. Martino, L., Brown, N. R., Masino, L., Esposito, D. & Rittinger, K. Determinants of E2-ubiquitin conjugate recognition by RBR E3 ligases. Sci Rep 8, (2018).

50. Yuan, L., Lv, Z., Atkison, J. H. & Olsen, S. K. Structural insights into the mechanism and E2 specificity of the RBR E3 ubiquitin ligase HHARI. Nat Commun 8, (2017).

51. Platanias, L. C. Mechanisms of type-I- and type-II-interferon-mediated signalling. Nature Reviews Immunology vol. 5 375–386 Preprint at 10.1038/nri1604 (2005).

52. Ma, F. et al. Positive feedback regulation of type I interferon by the interferon-stimulated gene STING. EMBO Rep 16, 202–212 (2015).

53. Gough, D. J., Messina, N. L., Clarke, C. J. P., Johnstone, R. W. & Levy, D. E. Constitutive Type I Interferon Modulates Homeostatic Balance through Tonic Signaling. Immunity vol. 36 166–174 Preprint at 10.1016/j.immuni.2012.01.011 (2012).

54. Richez, C., et al. Toll-like Receptor 4 Ligands Induce IFN-α Production By Mouse Conventional Dendritic Cells and Human Monocytes After IFN-β Priming 1 NIH Public Access. The Journal of Immunology J Immunol vol. 182 www.jimmunol.org. (2009).

55. Schweppe, D. K., Huttlin, E. L., Harper, J. W. & Gygi, S. P. BioPlex Display: An Interactive Suite for Large-Scale AP-MS Protein-Protein Interaction Data. J Proteome Res 17, 722–726 (2018).

56. Kleiger, G., Saha, A., Lewis, S., Kuhlman, B. & Deshaies, R. J. Rapid E2-E3 Assembly and Disassembly Enable Processive Ubiquitylation of Cullin-RING Ubiquitin Ligase Substrates. Cell 139, 957–968 (2009).

57. Crowe, C., et al. Mechanism of Degrader-Targeted Protein Ubiquitinability. Sci. Adv vol. 10 (2024).

58. Nguyen, D. M. et al. Structure and dynamics of a pentameric KCTD5/CUL3/Gß? E3 ubiquitin ligase complex. Proc Natl Acad Sci U S A 121, (2024).

59. Horn-Ghetko, D. et al. Noncanonical assembly, neddylation and chimeric cullin– RING/RBR ubiquitylation by the 1.8 MDa CUL9 E3 ligase complex. Nat Struct Mol Biol 31, 1083–1094 (2024).

60. Fu, D. J. & Wang, T. Targeting NEDD8-activating enzyme for cancer therapy: developments, clinical trials, challenges and future research directions. J Hematol Oncol 16, (2023).

61. Crow, Y. J. & Stetson, D. B. The type I interferonopathies: 10 years on. Nat Rev Immunol 22, 471–483 (2022).

62. Crow, M. K. & Ronnblom, L. Type i interferons in host defence and inflammatory diseases. Lupus Science and Medicine vol. 6 Preprint at 10.1136/lupus-2019-000336 (2019).

63. Hijazi, M., Smith, R., Rajeeve, V., Bessant, C. & Cutillas, P. R. Reconstructing kinase network topologies from phosphoproteomics data reveals cancer-associated rewiring. Nat Biotechnol 38, 493–502 (2020).

64. Casado, P. et al. Kinase-Substrate Enrichment Analysis Provides Insights into the Heterogeneity of Signaling Pathway Activation in Leukemia Cells. Sci. Signal 26, (2013).

