## Supplement for "The acidic N-terminus of HHARI and neddylation are essential for the activation and maintenance of RIG-I-mediated type I interferon response"

**Supplementary Materials and Methods**

**Phosphoproteomic experiments**

Phosphoproteomics experiments were performed using mass spectrometry with some technical modifications as described.[1,2] In brief, frozen cell pellets were lysed in 8M urea buffer and supplemented with phosphatase inhibitors (10 mM $Na_3VO_4$, 100 mM β-glycerol phosphate and 25 mM $Na_2H_2P_2O_7$ (Sigma)). 110µg of proteins were reduced, alkylated, and digested into peptides using trypsin as previously described.[3,4] Phosphopeptides were desalted and enriched using the AssayMAP Bravo (Agilent Technologies) platform. For desalting, reverse phase S cartridges (Agilent, 5 µL bed volume) were primed with 250 µL 99.9% acetonitrile (ACN) and 0.1% trifluoroacetic acid (TFA) and equilibrated with 250 µL 0.1% TFA at a flow rate of 10 µL/min. The samples were loaded at 20 µL/min, followed by an internal cartridge wash with 0.1% TFA at a flow rate of 10 µL/min. Peptides were then eluted with 105 µL of 1M glycolic acid with 50% ACN, 5% TFA and this is the same buffer for subsequent phosphopeptide enrichment. Phosphopeptides were enriched using 5µl Assay MAP $TiO_2$ cartridges on the Assay MAP Bravo platform. The cartridges were primed with 100µl of 5% ammonia solution with 15% ACN at a flow rate of 300 µL/min and equilibrated with 50 µL loading buffer (1M glycolic acid with 80% ACN, 5% TFA) at 10 µL/min. Samples eluted from the desalting were loaded onto the cartridge at 3 µL/min. The cartridges were washed with 50 µL loading buffer and the phosphopeptides were eluted with 25 µL 5% ammonia solution with 15% ACN directly into 25 µL 10% formic acid. Phosphopeptides were lyophilised in a vacuum concentrator and stored at -80˚C. Dried phosphopeptides were dissolved in 0.1% TFA and analysed by nanoflow ultimate 3000 RSL nano instrument was coupled on-line to a Q Exactive plus mass spectrometer (ThermoFisher Scientific). Gradient elution was from 3% to 28% solvent B in 60 min at a flow rate 250 nL/min with solvent A being used to balance the mobile phase (buffer A was 0.1% 137 formic acid in water and B was 0.1% formic acid in acetonitrile). The spray voltage was 1.95 kV and the capillary temperature was set to 255 ºC. The Q-Exactive plus was operated in data dependent mode with one survey MS scan followed by 15 MS/MS scans. The full scans were acquired in the mass analyser at 375- 1500m/z with the resolution of 70 000, and the MS/MS scans were obtained with a resolution of 17 500.

MS raw files were converted into Mascot Generic Format using Mascot Distiller and searched against the SwissProt database (released February 2021) with restriction to human entries using the Mascot search daemon. Parameters include allowing mass windows of 10 ppm and 25 mmu for parent and fragment mass to charge values, respectively, and a maximum of 2 missed cleavages. Fixed modification of carbomidomethyl (C) were considered and variable modifications included in the search were oxidation of methionine, pyro-glu (N-term) and phosphorylation of serine, threonine and tyrosine. Phosphopeptide quantification was performed using in-house software Pescal as described before[3] where the data was normalised by the sum of all intensities from a sample and intensity values of zero (not detected) were substituted by minimum intensity values across all phosphopeptides in the same sample divided by 10. The data was further normalised by variance stabilisation normalisation using the Bioconductor package Vsn (version 3.62.0) for downstream statistics and analysis.[5] R (version 4.2) was used for statistical analysis and data visualisation. MatrixTests package (version 0.1.9) was used for t-test analysis. Boxplots were plotted using the gglot2 package (version 3.3.6) and the volcano plot was plotted using the easylabel package (version 0.2.5).

**Supplementary Table 1 – List of reagents used, including expression vectors and siRNA species.**

| Reagent | Identifier |
|---|---|
| ***Expression plasmids*** | |
| pcDNA3-HA-p48 (IRF9) | Addgene, 11614 |
| pcDNA5-FRT/TO-FLAG-RNF216 (TRIAD3) | MRC PPU Reagents & Services DU27006 |
| pcDNA5-FRT/TO-FLAG-RNF216 (TRIAD3) aa1-137 | Generated by site-directed mutagenesis |
| pcDNA3-HA2-ROC1 | Addgene, 19897 |
| pcDNA3-HA2-ROC1 D97A | Generated by site-directed mutagenesis |
| pcDNA3-myc3-CUL1 | Addgene, 19896 |
| pcDNA3-myc3-CUL2 | Addgene, 19892 |
| pcDNA3-myc3-CUL3 | Addgene, 19893 |
| pcDNA3-myc3-CUL4A | Addgene, 19951 |
| pcDNA3-myc3-CUL4B | Addgene, 19922 |
| pcDNA3-myc3-CUL5 | Addgene, 19895 |
| pcDNA3-DN-hCUL1-FLAG (aa1-452) | Addgene, 15818 |
| pcDNA3-DN-hCUL2-FLAG (aa1-427) | Addgene, 15819 |
| pcDNA3-DN-hCUL3-FLAG (aa1-418) | Addgene, 15820 |
| pcDNA3-DN-hCUL4A-FLAG (aa1-440) | Addgene, 15821 |
| pcDNA3-DN-hCUL4B-FLAG (aa1-594) | Addgene, 15822 |
| pcDNA3-DN-hCUL5-FLAG (aa1-441) | Addgene, 15823 |
| pCMV6-AC | Origene, PS100020 |
| pCMV6-AN-Myc | Origene, PS100003 |
| pCMV-HA-2-RIG-I | MRC PPU Reagents & Services DU32183 |
| pCMV-HA-2-RIG-I K172R | Generated by site-directed mutagenesis |
| pCMV-FLAG-TRIAD1 | MRC PPU Reagents & Services DU21693 |
| pCMV-FLAG-TRIAD1 aa1-60 | Generated by site-directed mutagenesis |
| pCMV6-UBE2L3 | Origene, RG208780 |
| pCMV6-UBE2L3 C86S | Generated by site-directed mutagenesis |
| pFLAG/CMV2-HHARI | Addgene, 17450 |
| pFLAG/CMV2-HHARI C357A | Generated by site-directed mutagenesis |

| | |
|---|---|
| pFLAG/CMV2-HHARI W386A | Generated by site-directed mutagenesis |
| pFLAG/CMV2-HHARI Y387A | Generated by site-directed mutagenesis |
| pFLAG/CMV2-HHARI S427A | Generated by site-directed mutagenesis |
| pFLAG/CMV2-HHARI S427D | Generated by site-directed mutagenesis |
| pFLAG/CMV2-HHARI S514A | Generated by site-directed mutagenesis |
| pFLAG/CMV2-HHARI S514A | Generated by site-directed mutagenesis |
| pFLAG/CMV2-HHARI 1-407 | Generated by site-directed mutagenesis |
| pFLAG/CMV2-HHARI 1-336 | Generated by site-directed mutagenesis |
| pFLAG/CMV2-HHARI 1-242 | Generated by site-directed mutagenesis |
| pFLAG/CMV2-HHARI 1-153 | Generated by site-directed mutagenesis |
| pFLAG/CMV2-HHARI 1-100 | Generated by site-directed mutagenesis |
| pCMV6-Myc-HHARI 95-557 | Generated by PCR cloning |
| pCMV6-Myc-HHARI 152-557 | Generated by PCR cloning |
| pRC/CMV-JAK1K908E-VSV | Addgene, 139357 |
| pRC/CMV-JAK1-VSV | Addgene, 139356 |
| pRC/CMV-TYK2K930R-VSV | Addgene, 139346 |
| pRC/CMV-TYK2-VSV | Addgene, 139344 |
| ***siRNA*** | |
| DDX58 (RIG-I)<br>S: CCAGAAUUAUCCCAACCGA[T][T]<br>AS: UCGGUUGGGAUAAUUCUGG[T][T] | ThermoFisher, s24144[6] |
| IFIH1 (MDA-5)<br>S: GUAACAUUGUUAUCCGUUA[T][T]<br>AS: UAACGGAUAACAAUGUUAC[A][T] | ThermoFisher, s34498[6] |
| IRF3<br>S: GGAAGACAUUCUGGAUGAG<br>AS: CUCAUCCAGAAUGUCUUCC[dT][dT] | Merck custom siRNA[7] |
| MAVS<br>S: CCACCUUGAUGCCUGUGAA<br>AS: UUCACAGGCAUCAAGGUGG[dT][dT] | Merck custom siRNA[8] |
| Non-targeting control<br>Sequence unavailable | Integrated DNA technologies, 51-01-14-04 |

**Supplementary Table 2 – list of antibodies used**

| Antibody | Identifier | Species | Application |
|---|---|---|---|
| Anti-goat HRP | Santa Cruz Technologies, Sc-2354 | Mouse | WB |
| Anti-mouse IgG HRP | BD Pharmingen, 554002 | Goat | WB |
| Anti-rabbit IgG HRP | CST, #7074 | Rabbit | WB |
| ARIH1 | Abcam, Ab3891 | Goat | WB |
| FLAG-tag | CST, #14793 | Rabbit | WB, IP |
| FLAG-tag | CST, #8146 | Mouse | WB, IP |
| HA-tag (F-7) | Santa cruz technologies, Sc-7392 | Mouse | WB, IP |
| HA-tag (C29F4) | CST, #3724 | Rabbit | WB |
| JAK1 (6G4) | CST, #3344 | Rabbit | WB |
| JAK2 (D2E12) | CST, #3230 | Rabbit | WB |
| JAK3 (D1H3) | CST, #8827 | Rabbit | WB |
| MAVS | CST, #3993 | Rabbit | WB |
| MAVS-Alexa Fluor 488 | Santa cruz technologies, sc-166583 AF488 | Mouse | Confocal |
| MDA-5 (D74E4) | CST, #5321 | Rabbit | WB |
| Myc-tag HRP | CST, #2040 | Mouse | WB |
| Myc-tag (71D10) | CST, #2278 | Rabbit | WB |
| Phospho-JAK1 (Y1034/1035) (D7N4Z) | CST, #74129 | Rabbit | WB |
| Phospho-JAK2 (Y1008) (D4A8) | CST, #8082 | Rabbit | WB |
| Phospho-JAK3 (Y980/981) (D44E3) | CST, #5031 | Rabbit | WB |
| Phospho-STAT1 (Y701) | CST, #7649 | Rabbit | WB |
| Phospho-STAT2 (Y690) | CST, #88410 | Rabbit | WB |
| Phospho-STAT3 (Y705) (D3A7) | CST, #9145 | Rabbit | WB |
| Phospho-STAT5 (Y694) [E208] | Abcam, Ab32364 | Rabbit | WB |

| | | | |
|---|---|---|---|
| Phospho-TBK1 (S172) (D52C2) | CST, #5483 | Rabbit | WB |
| Phospho-TYK2 (Y1054/1055) (D7T8A) | CST, #68790 | Rabbit | WB |
| PTP-1B | ProteinTech, 11334-1-AP | Rabbit | WB |
| RIG-I (D14G6) | CST, #3743 | Rabbit | WB |
| SH-PTP2 (B-1) | Santa Cruz Technologies, Sc-7384 | Mouse | WB |
| STING | CST, #13647 | Rabbit | WB |
| TBK1 (D1B4) | CST, #3504 | Rabbit | WB |
| TC-PTP | R&D Systems, MAB1930-SP | Mouse | WB |
| TYK2 (D4I5T) | CST, #14193 | Rabbit | WB |
| UBE2L3 | Abcam, Ab108936 | Rabbit | WB |
| USP-18 (D4E7) | CST, #4813 | Rabbit | WB |
| UBE2L6 | Abcam, Ab109086 | Rabbit | WB |
| GAPDH-HRP | BIORAD, MCA4739P | Mouse | WB |

### Supplementary Figure 1

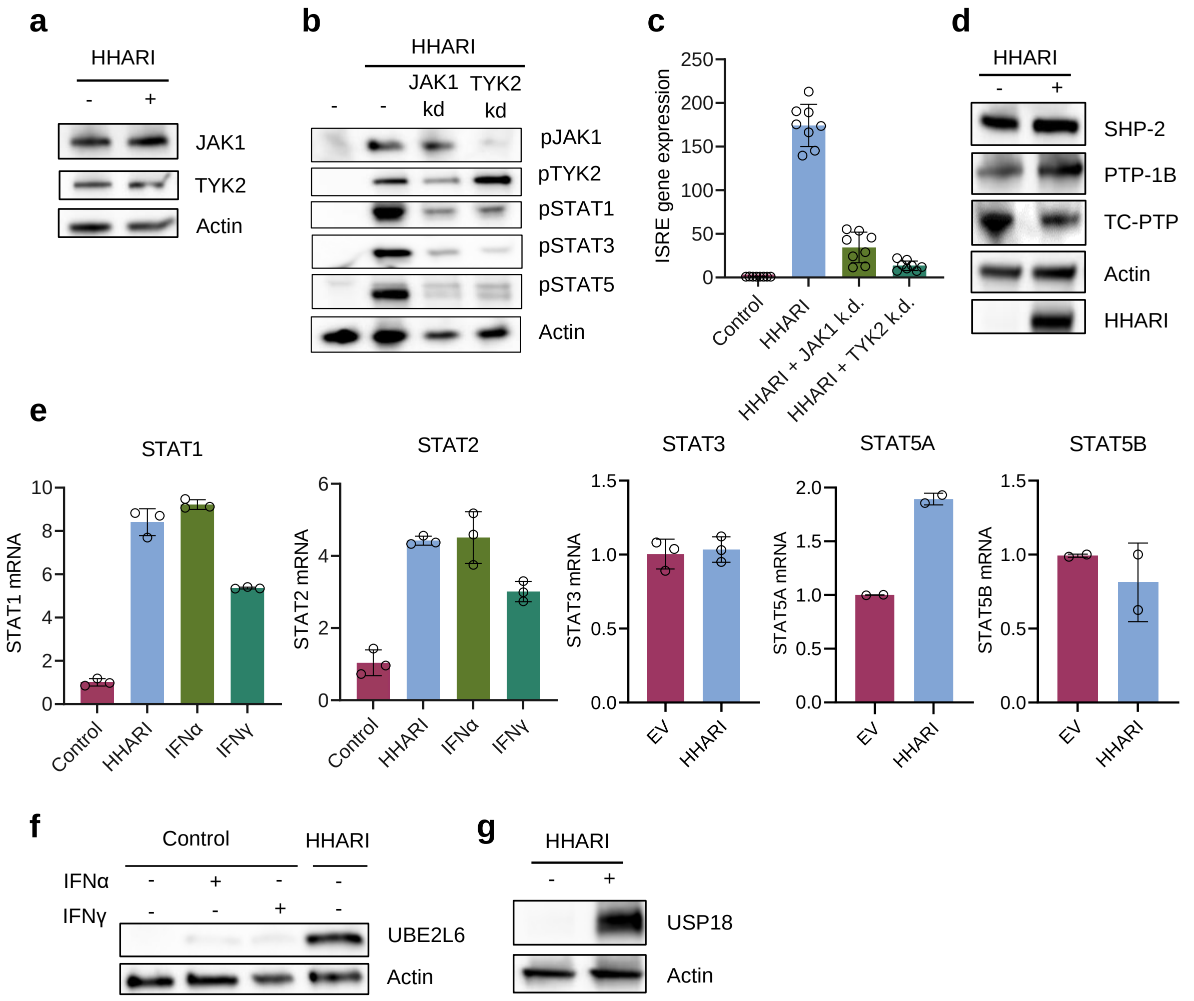


**Supplementary Figure 1 | HHARI overexpression leads to a type I IFN signature. (a)** Immunoblot of total JAK1, JAK2 and TYK2 protein levels in cells transfected with a control vector or FLAG-HHARI WT. **(b)** Immunoblot analysis of pJAK1, pTYK2, pSTAT1, pSTAT3 and pSTAT5 in cells transfected with FLAG-HHARI WT with/without kinase deficient JAK1 (K908E) or TYK2 (K930R) mutants. **(c)** ISRE luciferase reporter assay on cells treated as described under (b). **(d)** Immunoblot analysis of phosphatases including SHP-2, TC-PTP and PTP-1B in cells transfected with a control vector or FLAG-HHARI WT. **(e)** Fold changes in mRNA generated through real time reverse transcriptase PCR are shown. Genes investigated include *STAT1*, *STAT2*, *STAT3*, *STAT5A* and *STATB*. Cells were transfected with FLAG-HHARI or control vector for 48 hours. Cells were stimulated with 100ng/ml IFNα or IFNγ for 24 hours. Error bars represent standard deviation. **(f-g) U**BE2L6 and USP18 protein levels were analysed through immunoblot. Cells were transfected with FLAG-HHARI or control vector for 48 hours. Cells in (f) were also stimulated with 100ng/ml IFNα or IFNγ for 24 hours.

### Supplementary Figure 2

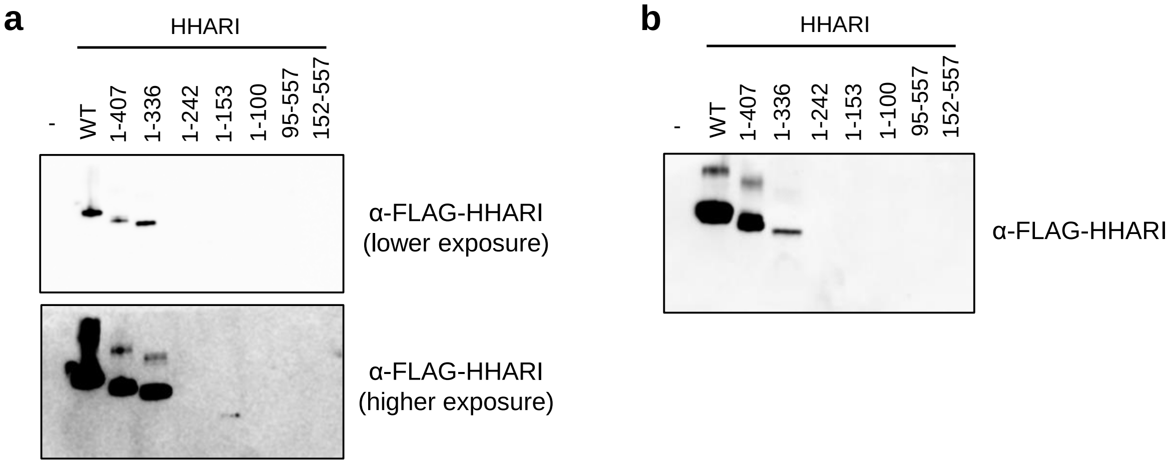


**Supplementary Figure 2 | Expression of HHARI truncation mutants varies. (a-b)** Immunoblot of FLAG-HHARI expression for FL (WT), 1-407, 1-336, 1-242, 1-153 and 1-100 mutants. Experiments a and b represent separate experiments. *Note: HHARI 95-557 and 152-557 constructs are myc-tagged and expression is shown in Figure 4d.*

**Supplementary References**


1. Casado P, Rodriguez-Prados JC, Cosulich SC, Guichard S, Vanhaesebroeck B, Joel S, et al. Kinase-Substrate Enrichment Analysis Provides Insights into the Heterogeneity of Signaling Pathway Activation in Leukemia Cells. *Sci Signal*. 2013;26(268). Available from: 10.1126/scisignal.2003573

2. Hijazi M, Smith R, Rajeeve V, Bessant C, Cutillas PR. Reconstructing kinase network topologies from phosphoproteomics data reveals cancer-associated rewiring. *Nat Biotechnol*. 2020;38(4):493–502.

3. Alcolea MP, Casado P, Rodríguez-Prados JC, Vanhaesebroeckp B, Cutillas PR. Phosphoproteomic analysis of leukemia cells under basal and drug-treated conditions identifies markers of kinase pathway activation and mechanisms of resistance. *Molecular and Cellular Proteomics*. 2012;11(8):453–66.

4. Montoya A, Beltran L, Casado P, Rodríguez-Prados JC, Cutillas PR. Characterization of a TiO2 enrichment method for label-free quantitative phosphoproteomics. *Methods*. 2011;54(4):370–8.

5. Huber W, von Heydebreck A, Sültmann H, Poustka A, Vingron M. Variance stabilization applied to microarray data calibration and to the quantification of differential expression. *Bioinformatics*. 2002;18(1):96–104.

6. Yoshino H, Iwabuchi M, Kazama Y, Furukawa M, Kashiwakura I. Effects of retinoic acid-inducible gene-i-like receptors activations and ionizing radiation cotreatment on cytotoxicity against human non-small cell lung cancer in vitro. *Oncol Lett*. 2018;15(4):4697–705.

7. Basit A, Cho MG, Kim EY, Kwon D, Kang SJ, Lee JH. The cGAS/STING/TBK1/IRF3 innate immunity pathway maintains chromosomal stability through regulation of p21 levels. *Exp Mol Med*. 2020;52(4):643–57.

8. Seth RB, Sun L, Ea CK, Chen ZJ. Identification and characterization of MAVS, a mitochondrial antiviral signaling protein that activates NF-κB and IRF3. *Cell*. 2005;122(5):669–82.
